# Systematic gene tagging using CRISPR/Cas9 in human stem cells to illuminate cell organization

**DOI:** 10.1101/123042

**Authors:** Brock Roberts, Amanda Haupt, Andrew Tucker, Tanya Grancharova, Joy Arakaki, Margaret A. Fuqua, Angelique Nelson, Caroline Hookway, Susan A. Ludmann, Irina A. Mueller, Ruian Yang, Alan R. Horwitz, Susanne M. Rafelski, Ruwanthi N. Gunawardane

## Abstract

We present a CRISPR/Cas9 genome editing strategy to systematically tag endogenous proteins with fluorescent tags in human inducible pluripotent stem cells. To date we have generated multiple human iPSC lines with GFP tags for 10 proteins representing key cellular structures. The tagged proteins include alpha tubulin, beta actin, desmoplakin, fibrillarin, lamin B1, non-muscle myosin heavy chain IIB, paxillin, Sec61 beta, tight junction protein ZO1, and Tom20. Our genome editing methodology using Cas9 ribonuclear protein electroporation and fluorescence-based enrichment of edited cells resulted in <0.1-24% HDR across all experiments. Clones were generated from each edited population and screened for precise editing. ∼25% of the clones contained precise mono-allelic edits at the targeted locus. Furthermore, 92% (36/39) of expanded clonal lines satisfied key quality control criteria including genomic stability, appropriate expression and localization of the tagged protein, and pluripotency. Final clonal lines corresponding to each of the 10 cellular structures are now available to the research community. The data described here, including our editing protocol, genetic screening, quality control assays, and imaging observations, can serve as an initial resource for genome editing in cell biology and stem cell research.

## Introduction

The study of cellular processes using new genome editing strategies, particularly CRISPR/Cas9, is becoming increasingly feasible and powerful [1-5]. Several experimental strategies to harness the utility of CRISPR/Cas9-mediated genome editing is in use with the most common being gene disruption for loss of function analysis. This approach exploits the error-prone non-homologous end joining (NHEJ) pathway of DNA repair [6]. CRISPR/Cas9 is also used to introduce polymorphisms, often associated with disease, via the much less efficient homology directed repair (HDR) pathway [7].

One of the more challenging applications of CRISPR/Cas9 is the introduction of large tags, such as GFP, into the genome. With this capability, one can precisely target a genomic locus, inserting a fluorescent protein under the endogenous regulatory control of the native protein [3]. Inserting a large tag sequence requires that the cell utilizes HDR with an exogenously provided repair template, a process that is inefficient in human cells. Nevertheless, the ability to endogenously tag proteins in cells offers a major improvement over conventional overexpression systems, especially in live imaging and functional studies [3, 8, 9].

While a growing number of studies illustrate the power of endogenous gene tagging, it has only been used comprehensively in budding yeast [10, 11]. Strategies are now emerging to tag multiple genes in standard human cell line models [12, 13]. They all employ different approaches using either selection or extensive screening to overcome the inherent inefficiency of HDR [7]. Due to the inefficiency of this process, it is unclear whether HDR-mediated tagging is sufficiently reproducible and precise across targets and cellular contexts to enable its systematic use. To this end, we have generated a collection of human induced pluripotent stem cell (hiPSC) lines in which the GFP sequence is introduced at multiple genomic loci encoding various cellular proteins [14]. We chose proteins that localize to key cellular structures for our tagging experiments. The resulting endogenously GFP-tagged cell lines can be used for live cell imaging to elucidate the organization and dynamics of the major organelles and structures of the cell, providing an important resource to study human cell biology.

We selected hiPSCs for this genome editing effort for several reasons [15-17]. Since hiPSCs are derived from individuals with characterized physiology, they are relevant to human disease, providing “disease in a dish” models [18-20]. Furthermore, hiPSCs provide the opportunity to study the tagged proteins in a diploid and non-transformed cellular context. Because hiPSCs can be differentiated into multiple cell types, they also offer the opportunity to study tagged gene products in multiple cellular contexts. Finally, the propensity of hiPSCs to maintain a stable karyotype over dozens of passages in culture enables the genome editing and clonal line generation processes. We chose the Wild Type C (WTC) hiPSC line derived from a healthy donor as our parental line for all gene edits for multiple reasons including its episomal derivation, known genomic stability, availability of genomic sequence, differentiation into diverse cell fates, and open access to the academic research community [21, 22].

Here we present our methodology for CRISPR/Cas9-mediated genome editing to create a collection of isogenic, clonally derived WTC hiPSC lines labeling 10 key intracellular structures, and the genetic screening and quality control data accompanying this effort. With this method, we find that **1)** GFP tag knock-in was successful for 10 targeted genes from diverse loci with typical editing efficiencies ranging from 0.1-4%; **2)** precisely edited clones were identifiable with PCR-based analyses for 9 of these 10 experiments; **3)** 92% (36/39) of the final candidate clonal lines generated from these editing experiments met all quality control criteria, including correct expression and subcellular localization of the tagged protein, normal karyotype and morphology, and robust pluripotency including multi-lineage differentiation potential. To our knowledge, this is the first systematic study that introduces, characterizes and validates genome-edited hiPSCs expressing complete GFP sequences. Additionally, our data illustrate that systematic GFP-tagging of diverse loci in hiPSCs is feasible and can generate high quality stem cell lines for diverse applications.

## Results

### Genome editing strategy

We used CRISPR/Cas9 and HDR to incorporate a full length GFP tag sequence into 10 genomic loci to label a group of key intracellular structures (Table 1). Experiments were designed to introduce the GFP tag at the N- or C-terminus along with a short peptide linker between the endogenous protein and the GFP tag (Table 1). The strategy used for N-terminal tagging is shown as an example (Fig. 1A). We confirmed expression of the transcript isoform(s) designated for tagging by performing RNA-Seq on the parental WTC line prior to genome editing (Fig. S1 and data not shown) [23-25].

**Figure 1.**
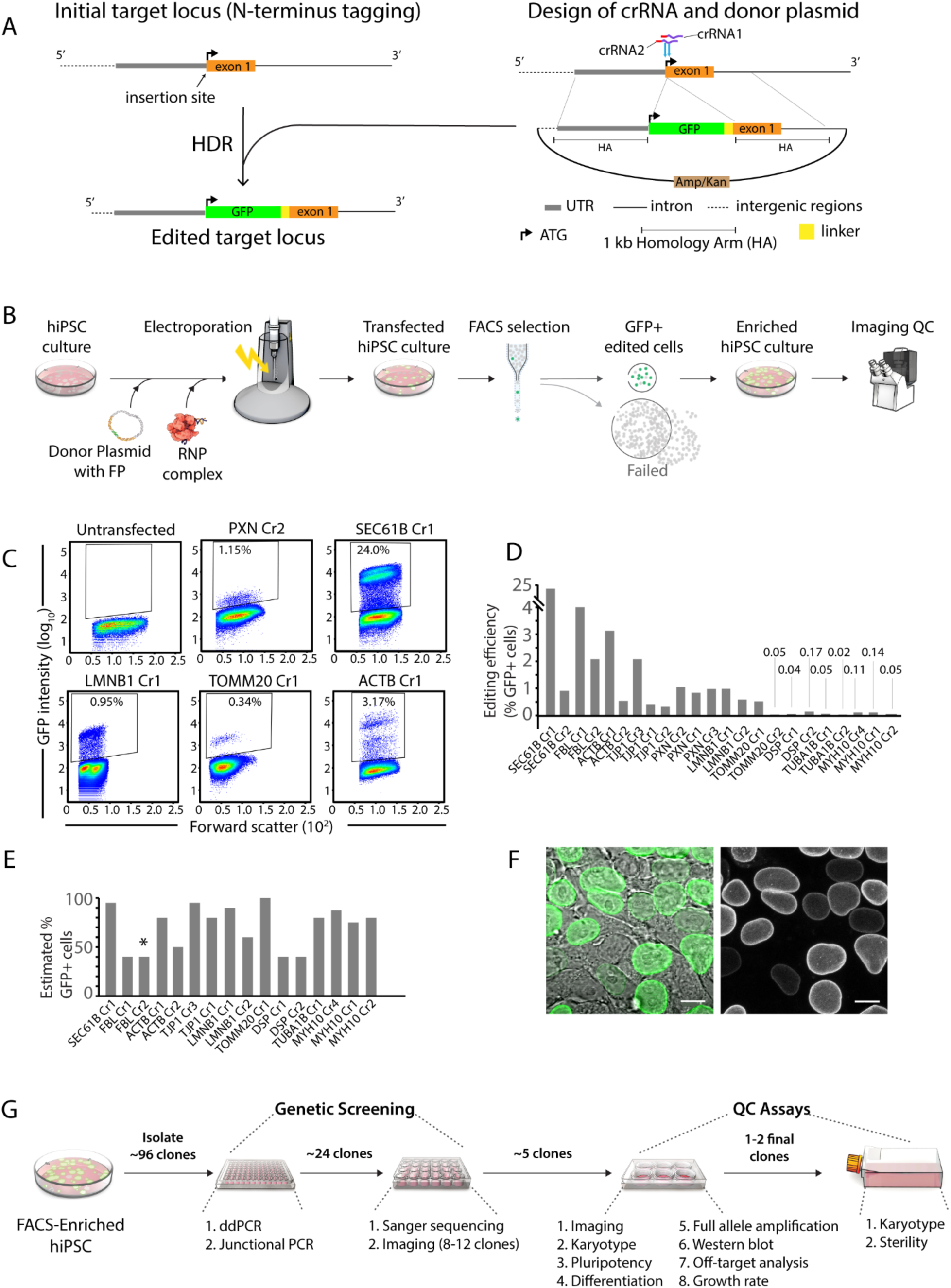
Genome editing experimental design and initial comparisons of editing efficiency. **(A)** Schematic illustrating design features important for genome editing experiments. An N-terminal GFP tagging strategy preceding the first exon of the gene of interest is shown as an example. The location of both crRNA binding sequences is indicated in purple. PAM sequences are indicated in red. The position of the anticipated double strand break generated by each crRNA is indicated with a blue arrow. The donor plasmid contains 1 kb homology arms on either side of the GFP and linker sequence and a bacterial selection sequence in the backbone. The example in the schematic shows successful N-terminal tagging via HDR resulting in the tag and linker being inserted after the endogenous start codon (ATG) in frame with the first exon. **(B)** Schematic depicting the genome editing process. Transfection includes pre-complexing of the RNP (Cas9/crRNA/tracrRNA) and co-electroporation with the donor plasmid. FACS is used to enrich for GFP-positive cells 3-4 days after transfection. GFP-positive cells are collected and expanded as an enriched population for image-based confirmation of tagging before clonal line generation. **(C)** Flow cytometry plots displaying GFP intensity (y-axis) 3-4 days after editing. Data shown from several experiments (target locus is indicated), along with control untransfected cells. Gates indicate the population of putatively edited cells and values reflect the percentage of edited cells within the total population. Forward scatter is shown on the x-axis. **(D)** Comparison of genome editing efficiency, as defined by FACS, shown as a percentage of GFP-positive cells within the gated cell population in each experiment. **(E)** Estimated percentage of cells in the FACS-enriched populations expressing GFP, as determined by live microscopy. This analysis was not performed on PXN edited cells. The majority (>50%) of GFP-positive cells in each case displayed correctly localized GFP, except where indicated by an asterisk; only ∼5% of GFP-positive cells in the FBL Cr2 population had correct subcellular localization. **(F)** Representative image of the LMNB1 Cr1 FACS-enriched population showing an enrichment of GFP-positive cells. As expected, the edited population is a mixture of GFP-positive and GFP-negative cells. GFP intensity level is also variable. Scale bars are 10 μm. **(G)** Schematic overview of the clone isolation, genetic screening, and quality control workflow. The genetic screening and quality control assays help identify 1-2 final clones from each gene tagging experiment.

**Table 1.** Summary of tagged structures.

| Gene | Protein | Cellular structure | Terminus tagged | Linker | Linker-specific reference |
| --- | --- | --- | --- | --- | --- |
| PXN | Paxillin | Matrix adhesions | C-terminus | GTSGGS | (commonly used linker sequence) |
| TUBA1B | Alpha tubulin | Microtubules | N-terminus | GGSGGS | Gan, et al. 2016 |
| DSP | Desmoplakin | Desmosomes | C-terminus | HDPPVAT | Godsel, et al. 2005 |
| LMNB1 | Nuclear lamin B1 | Nuclear envelope | N-terminus | SGLRSRAQAS | Michael Davidson Fluorescent Protein Collection |
| TOMM20 | Tom20 | Mitochondria | C-terminus | GGSGDPPVAT | Michael Davidson Fluorescent Protein Collection |
| ACTB | Beta actin | Actin filaments | N-terminus | AGSGT | Grassart, et al. 2014 |
| SEC61B | Sec61 beta | Endoplasmic reticulum | N-terminus | SGLRS | Shibata, et al. 2008 |
| FBL | Fibrillarin | Nucleolus | C-terminus | KPNSAVDGTAGPG | Dundr, et al. 2000 |
| MYH10 | Non-muscle myosin heavy chain IIB | Actomyosin bundles | N-terminus | YSDLELKLRIP | Wei and Adelstein 2000 |
| TJP1 | Tight junction protein ZO1 | Tight junctions | N-terminus | SGLRSRALERDK | Riesen, et al. 2002 |

The 10 genes and corresponding proteins successfully targeted in genome editing experiments are listed. The peptide terminus chosen for introduction of the GFP tag, and the peptide linker chosen to fuse the GFP tag with the endogenous protein are shown.

We utilized the ribonuclear protein (RNP) method with recombinant wild type *S. pyogenes* Cas9 (SpCas9) protein pre-complexed with a synthetic CRISPR RNA (crRNA) and trans-activating crRNA (tracrRNA) duplex [26, 27]. This RNP complex was co-electroporated into the WTC cells along with GFP donor plasmids (Fig. 1B). The donor plasmids contained 1kb homology arms (HAs) and a target-specific linker sequence (Fig. 1A, Table 1) [28-34]. Other design features specific to each target locus were included the donor plasmid homology arms, including single nucleotide polymorphisms (SNPs) and insertions or deletions (INDELs) specific to the WTC genome and mutations to inactivate crRNA binding sites that would otherwise remain intact after the introduction of the linker and GFP sequence (Table S1) [22]. At least two independent crRNA sequences were used in each editing experiment to maximize editing success (Fig. 1A, Table S1). When possible, we only used crRNAs targeting Cas9 to within 50 bp of the intended GFP integration site with a strong preference for any crRNAs with binding sites within 10 bp (Table S1)[35, 36]. Cas-OFFinder was used to select and rank available crRNA sequences with respect to their genome-wide specificity (Fig. S2) [37]. Only crRNAs unique within the human genome were used with one unavoidable exception (TOMM20), and crRNAs whose alternative binding sites include mismatches in the “seed” region and are in non-genic regions were prioritized for use whenever possible (Fig. S2) [38-40].

After co-transfection of the RNP and donor plasmid, cells were recovered, sorted and analyzed for GFP expression by fluorescence assisted cell sorting (FACS) (Fig. 1B, C, S3). We observed GFP fluorescence in putatively edited cells at variable rates over a range of signal intensities (Fig. 1C, S3). We used the percentage of GFP-positive cells above the background defined by untransfected cells as a measure of HDR-mediated knock-in efficiency (Fig. 1D). Successful GFP-tagging was observed using at least one crRNA at 10 of the 12 loci even when HDR was inefficient (<1%). Across the 10 targets resulting in successful editing, the efficiency of HDR ranged from <0.1%-4%, with SEC61B a notable exception at 24% (Fig. 1D). In many cases, we observed that HDR efficiency at a given locus depended greatly on the crRNA used.

As expected for tagging experiments targeting diverse cellular proteins, the observed GFP intensity among GFP-positive cells varied widely. We observed weak GFP signal in some experiments where the target gene transcript was relatively scarce (PXN) and/or the protein localizes to distinct, discreet puncta (DSP) (Fig. 1C, S1, S3, 3C, 3F). We were nevertheless able to enrich for cells edited at these loci, although their expression level relative to the background threshold of unedited cells was minimal (Fig. 1C, S3). The two cases of failure were GALT, a relatively rare transcript, and TUBG1, whose product is known to localize to discreet puncta that may have insufficient signal for flow based detection (Fig. S1, S3) [41, 42]. In cases where we observed cells with varying GFP intensities (Fig. 1C, S3, e.g. SEC61B, LMNB1, ACTB), all GFP-positive cells were sorted as a single population (“FACS-enriched population”) for further characterization and clone generation.

Live imaging of the FACS-enriched populations was performed to assess the percentage of GFP-positive cells after sorting and confirm GFP localization to the appropriate structure in at least some of the cells. This served as an important validation step for precise editing prior to clone generation. We observed GFP-positive cells in 40-100% of the imaged cells across editing experiments (Fig. 1E). These enriched populations were generally mixed with respect to the fraction of GFP-positive cells and levels of GFP intensity (Fig. 1E, F). In all experiments except FBL Cr2 (where only ∼5% of GFP-positive cells had anticipated nucleolar GFP localization), the majority of GFP-positive cells displayed GFP localization to the appropriate cellular structure (Fig. 1F, and data not shown). We hypothesize that the variance in the localization (e.g. FBL) and intensity of the GFP signal (e.g. LMNB1), where observed, may reflect heterogeneous genome editing outcomes, with some cells incorporating the GFP tag imprecisely.

We subsequently generated clonal lines starting from these edited, enriched cell populations to identify and isolate precisely edited cells. Since stem cells are known to be sensitive to single cell isolation, we seeded cells at clonal density and manually isolated ∼10^2^ colonies per target gene. Using our isolation method, we achieved 79% clone survival across all experiments (Table S2). Following clonal recovery, we subjected these clones to genetic screening, as described below, followed by a suite of quality control assays to select our final, high quality clones based on stringent genomic, phenotypic, cell biological and stem cell criteria (Fig. 1G).

### Genetic screening of clones

We designed several PCR assays to screen for clones with precise editing. Our criteria for precise editing was incorporation of the GFP tag in frame with the targeted exon at the intended locus, the absence of donor plasmid backbone integration, and no mutations in either allele upon sequencing. Towards this aim, we employed a 3-step PCR screening approach (Fig. 2A). The first step was to identify GFP-tagged clones with no plasmid backbone integration using droplet digital PCR (ddPCR) (Fig. 2A left panel, step 1). We first quantified the abundance of the GFP tag sequence (x-axis in Fig. 2A left panel, 2B) and normalized this value to a known 2 copy genomic reference gene (RPP30) in order to calculate genomic GFP copy number in the sample.

**Figure 2.**
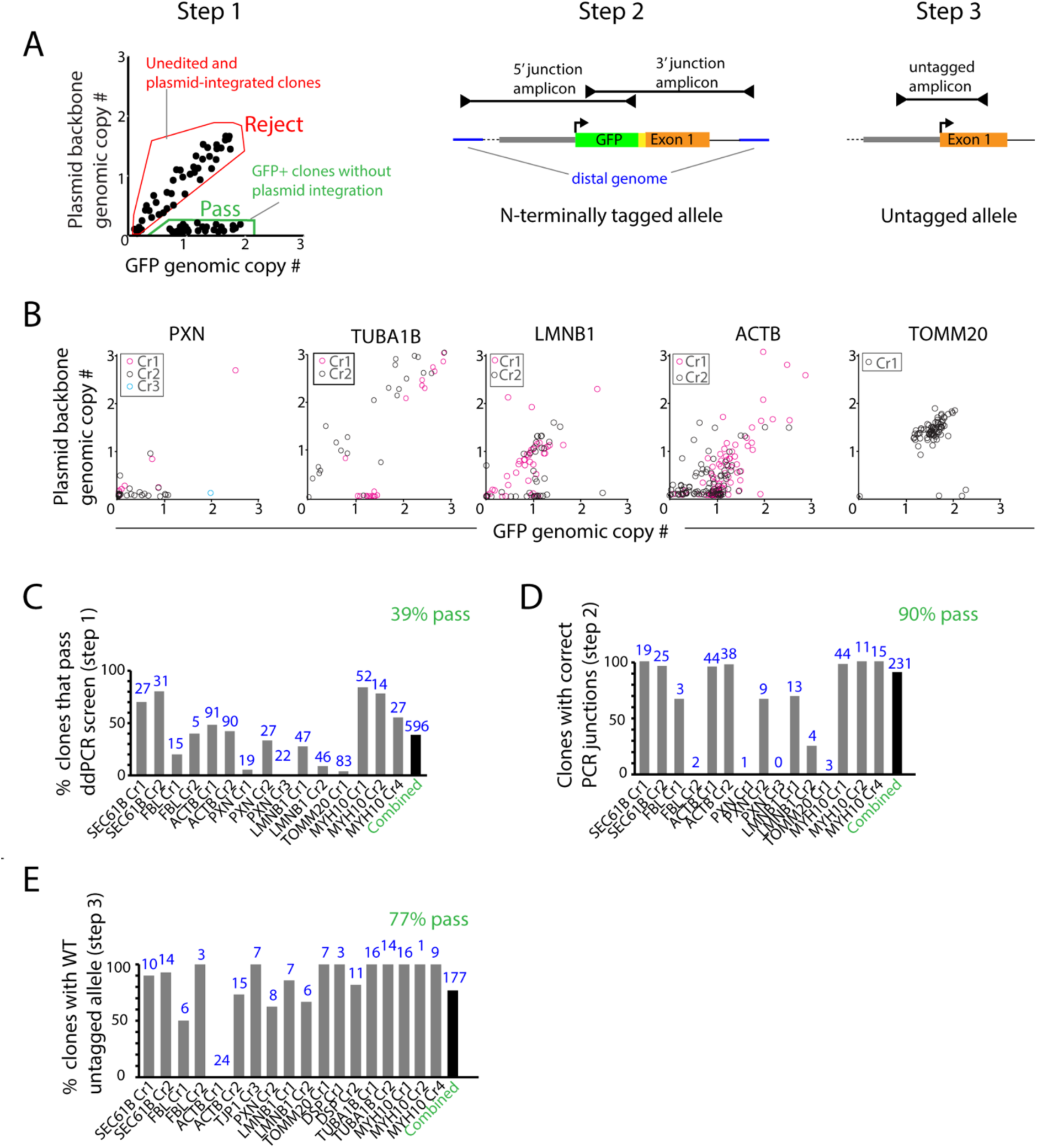
Genetic assays to screen for precise genome editing in clones. **(A)** Schematic illustrating the sequential process for identifying precisely tagged clones. In step 1 (left panel), ddPCR is used to identify clones with GFP insertion (normalized genomic GFP copy number ∼1 or ∼2) and no plasmid integration (normalized genomic plasmid backbone copy number <0.2). Hypothetical example of a typical editing experiment is shown with examples for pass (green) and fail (red) criteria. In step 2 (middle panel), junctional PCR amplification of the tagged allele is used to determine precise on-target GFP insertion. In step 3 (right panel), the untagged allele of a clone with mono-allelic GFP insertion is amplified. The amplicon is then sequenced to ensure that no mutations have been introduced to this allele. **(B)** Examples of ddPCR screening data (step 1) from experiments representative of the range of outcomes observed. Each data point represents one clone. Clones with GFP genomic copy number of ∼1 to ∼2 and plasmid backbone genomic copy number <0.2 were considered for further analysis. (**C**) Step1 results: percentage of clones confirmed by ddPCR to have incorporated the GFP tag but not the plasmid backbone. Data is shown across experiments where ddPCR was performed as the initial screen. (**D**) Step 2 results: percentage of clones confirmed in step 1 that also have correctly sized junctional PCR amplicons. (**E**) Step 3 results: percentage of clones confirmed to have wild type untagged alleles by PCR amplification and Sanger sequencing following steps 1 and 2. (**C-E**) Number of clones analyzed for each experiment is shown in blue. Percentage of clones across all experiments (combined) that met the screening criteria (C-E) is shown in green (39%, 90%, and 77%). TUBA1B, DSP and TJP1 analysis is omitted from C and D because junctional screening (step 2) was performed before ddPCR screening (step 1) in these experiments.

As part of the same analysis step we calculated the copy number of the donor plasmid (using the AMP or KAN sequence) in each clone (y-axis in Fig. 2A, left panel, 2B). We used a relatively conservative copy number threshold of 0.2 copies/genome as the cut off for eliminating clones with integrated plasmid. Overall, we rejected 45% of clones (Fig. S6), combining data across all 10 successful editing experiments with this criterion. Clones with a GFP copy number of ∼1.0 or ∼2.0 and AMP/KAN <0.2 were putatively identified as correctly edited mono- or bi-allelic GFP clones. Clones with GFP copy number 0.2-1 were considered mosaics of edited and unedited cells and were typically rejected (Fig. 2A, B, S4). The abundance of unedited and mosaic clones (0.2-1 GFP copies) observed for target genes such as PXN reflect the relative difficulty of enriching for endogenously tagged proteins with low expression (Fig. 1C, 2B and S1).

Since the probes for GFP, donor plasmid, and the RPP30 reference gene could be applied to all gene edits, this multiplexed ddPCR assay was used to rapidly interrogate large sets of clones in parallel without having to optimize parameters specifically for each target gene [7, 43, 44]. Overall, 39% (n=596) of the clones tested in step 1 satisfied our dual criteria of GFP incorporation with no evidence of plasmid integration (Fig. 2C). The rate of precise editing in step 1 varied based both on the locus and the crRNA used. The clones that passed step 1 were almost exclusively mono-allelic (Fig. 2B, C). Putatively bi-allelic GFP clones with no plasmid incorporation were rare (Fig. 2B). Therefore, we further screened clones with a GFP copy number between ∼1 and ∼2 to potentially identify bi-allelic clones from mixed cultures.

However, the majority of these clones (6 of 8) showed evidence of faulty DNA repair in the subsequent analysis step as discussed below (Fig. 2B, S4).

After confirmation of GFP tagging and the absence of plasmid integration in step 1, we used junctional PCR to determine whether the GFP tag was precisely incorporated without large duplications or deletions, common known byproducts of HDR (Fig. 2A – middle panel, step 2)[45, 46]. We amplified two overlapping amplicons that spanned the 5′ and 3′ junctions between the GFP tag and the host cell genome distal to the 1 kb homology arm sequences present in the donor plasmid (Fig. 2A, step 2). While a high rate of correct junctional products was observed for most gene edits, certain loci were more error prone (e.g. PXN Cr1 and TOMM20 Cr1). The small number of clones analyzed (passing step 1) in these cases may also indicate locus-specific challenges for precise editing. Notably, 6 of 8 putative bi-allelic clones discovered in several experiments failed this junctional PCR step. Nevertheless, 90% (n=231) of the overall clones tested in this second step contained expected junctional PCR products (Fig. 2D). Sanger sequencing the junctional amplicons from a subset of these clones (n=107) confirmed correct sequences in all clones (data not shown). The high rate of success in this assay validated the use of ddPCR to screen out clones with imprecise HDR.

In a third and final step, the untagged allele (for mono-allelic GFP-tagged clones) was amplified and sequenced in order to ensure that no mutations had been introduced via the alternative and preferred NHEJ pathway at the binding site of the crRNA used for editing (Fig. 2A, step 3, 2E). We chose a subset of clones confirmed by ddPCR and junctional PCR from each gene edit and performed Sanger sequencing on the amplification product corresponding to the untagged allele. Clones in which the untagged allele had become mutated via error-prone NHEJ were rejected. 77% (n=177) of the clones analyzed from all experiments contained a wild type untagged allele (Fig. 2E). A subset of these clones was chosen for further analysis in quality control assays (Fig. 1G).

In follow-up assays on the final candidate clones, we confirmed the presence of both the tagged and untagged alleles in a single, non-tiled junctional PCR reaction to rule out the possibility of misleading junctional PCR outcomes such as rearrangements and duplications (Fig. S5). In 9 out of 10 cases, we confirmed the presence of the expected products for both the tagged and untagged alleles (Fig. S5). TOMM20 was the exception, as discussed below. In the absence of precise editing at this locus, we chose several TOMM20 clones with evidence of plasmid backbone insertion in the non-coding sequences at the TOMM20 locus for further analysis.

Taken together, we observed overall confirmation rates of 39% (GFP incorporation with no plasmid), 90% (correct junctions), and 77% (wild type untagged allele) in each of the three screening steps across all gene targeting experiments (Fig. 2C-E). Only ∼25% of the clones screened in this manner met all three of these precise editing criteria. Donor plasmid integration was the most common category of imprecise editing, affecting 45% of all clones (Fig. S6). Our data suggest that this frequently occurs at the edited locus as a faulty byproduct of the editing process and that screening by junctional PCR alone would have led to clones with imprecise editing (Fig. S5, S6) [45, 46].

Locus- and crRNA-specific, widespread faulty editing was observed in some experiments. For example, TOMM20 yielded GFP-positive cells from only one crRNA (Cr1), all of which contained integrated plasmid (80/83) or faulty junctions (3/83) (Fig. 2 B-D, S3, S5, S6). The large majority of TUBA1B clones edited with Cr2 contained integrated plasmid, while most clones from Cr1 were unaffected. Similarly, the frequency and type of mutations found in the unedited allele were also target and locus specific, with ACTB Cr1 a notable outlier case in which NHEJ-mediated mutations in the untagged allele occurred in all analyzed clones (n=24) (Fig. 2E).

Clones with ddPCR signatures consistent with bi-allelic editing (GFP copy number ∼2) were observed at low frequency across all experiments (total n=8) (Fig. 2B, S4). Only one clone (PXN Cr2 cl. 53) was confirmed as a bi-allelic edit with predicted junctional products (Fig. 2A, step 2) and absence of the untagged allele (Fig. 2A, step 3) (data not shown), but was later rejected due to poor morphology (Fig. 5A). Other suspected bi-allelic clones were rejected due to incorrect junctional products (Fig. 2A, step 2) and/or presence of the untagged allele (Fig. 2A, step 3) indicating that these clones did not precisely incorporate the GFP tag in both alleles. The frequency of faulty HDR demonstrated by these data underscores the importance of our multistep genomic screening approach to identify precisely edited clones and confirm mono- and biallelic editing.

**Figure 5.**
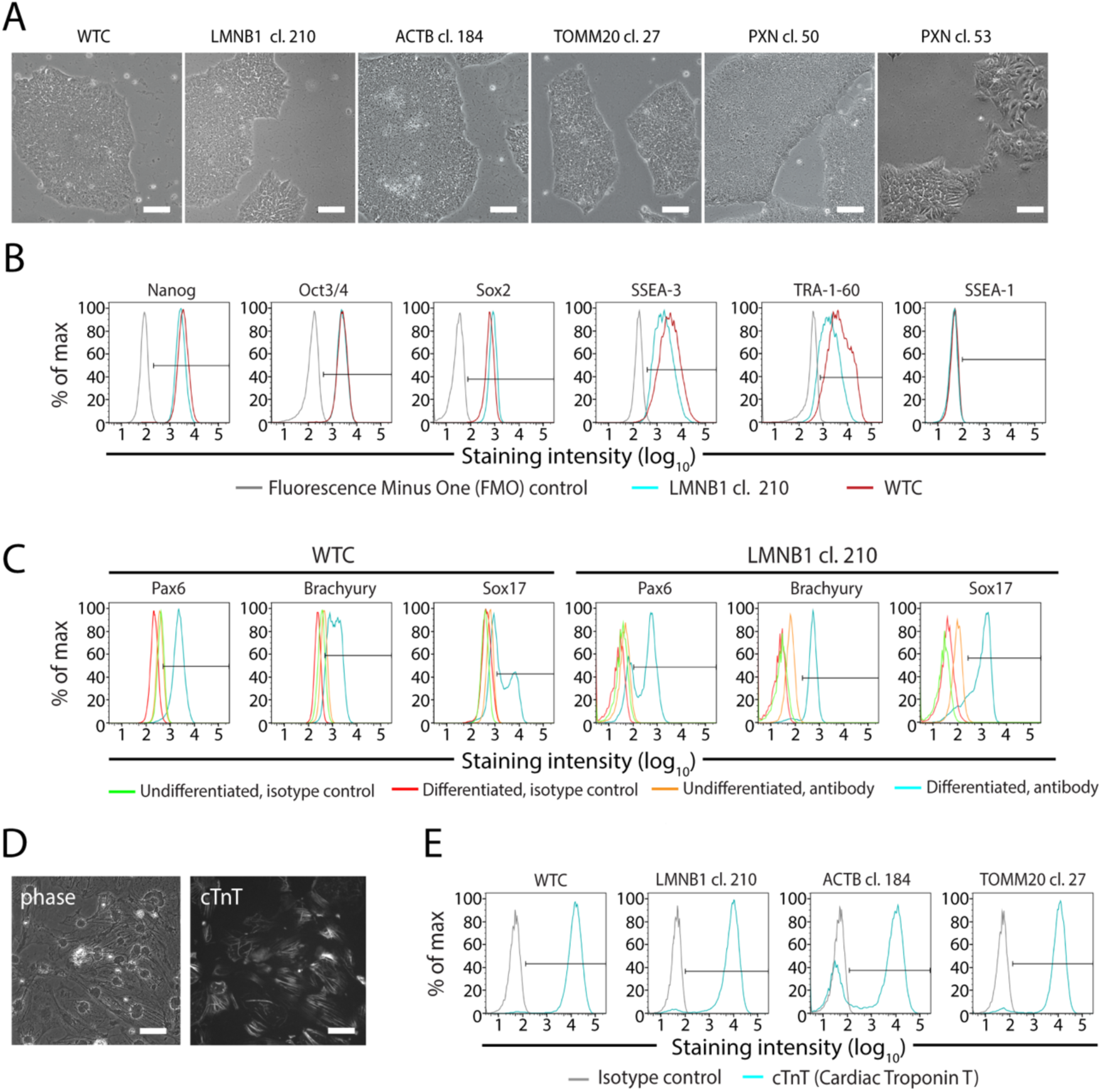
Assessment of stem cell quality after genome editing. (**A**) Representative phase contrast images depicting cellular and colony morphology of the unedited WTC line and several GFP-tagged clones (LMNB1, ACTB, TOMM20, and PXN). Images are of mature stem cell colonies after expansion. PXN cl. 53, rejected due to poor stem cell morphology, is counterexample. Scale bars are 100 μm. (**B**) Representative flow cytometry plots of gene-edited LMNB1 cl. 210 cells (blue), and unedited WTC cells (red) immunostained for indicated pluripotency markers (Nanog, Oct3/4, Sox2, SSEA-3, TRA-1-60) and a marker of differentiation (SSEA-1). Fluorescence Minus One (FMO) controls (grey) defined the positive staining threshold. (**C**) Representative flow cytometry plots of differentiated unedited WTC cells or gene-edited LMNB1 cl. 210 cells immunostained for markers of ectoderm (Pax6), mesoderm (Brachyury), and endoderm (Sox17) lineages. Differentiated cells stained with isotype control antibody (red) were used to define the positive staining threshold. Undifferentiated, gene-edited or WTC unedited cells (yellow) and their respective isotype controls (green) are overlaid. (**D**) Cardiomyocytes differentiated from unedited WTC cells and stained with cardiac Troponin T (cTnT) antibody to label cardiac myofibrils. Scale bars are 50 μm. (**E**) Representative flow cytometry plots showing cTnT expression (blue) in unedited WTC control cells and several gene edited cell lines (LMNB1 cl. 210, ACTB cl. 184, and TOMM20 cl. 27). Isotype controls (grey) defined the positive staining threshold. (**B-E**) Antibody information available in Table S4.

To assess whether the clones that met the above gene editing criteria contained off-target mutations due to non-specific CRISPR/Cas9 activity, we analyzed several final candidate clones from each experiment for mutations at off-target sites predicted by Cas-OFFinder (Fig. S2, S7)[37]. Potential off-target sites for each crRNA were prioritized for screening based both on their similarity to the on-target site used for editing and their proximity to genic regions (see methods). PCR amplification of these regions followed by Sanger sequencing was performed to identify potential mutations in 3-5 final candidate clones for all 10 genome editing experiments (6-12 sequenced sites per clone) across 142 unique sites. We were unable to identify any off-target editing events among a total of 406 sequenced loci (Fig. S7).

Establishing clonal hiPSC lines and culturing them long term is known to carry the risk of fixing somatic mutations and introducing genomic instability. To address this concern in our genome-edited lines, we karyotyped each candidate clone during our clonal line generation and expansion process (Fig. 1G). Of the 39 final candidate clonal lines tested, we only detected a karyotypic abnormality in one candidate clone (TUBA1B), which was then rejected (data not shown). Therefore, our data indicate that chromosomal abnormalities arise at a permissively low rate for high-throughput editing in the WTC line using our methodology.

Our assays described above allowed us to identify and expand a refined set of 5-10 candidate clones for further analysis of genomic, cell biological, and stem cell integrity as described below (Fig. 1G). These quality control assays were often performed in sequence, with only a few final clones being tested in all assays (Fig. 1G, Table 4). Although multiple crRNAs were tested in parallel in each editing experiment, only one crRNA per target locus generated clones under consideration for the final clonal line in 8/10 experiments (DSP and SEC61B were the exceptions). The remaining crRNAs resulted in inefficient or imprecise HDR (TOMM20, TJP1, TUBA1B, ACTB, PXN), altered morphology and/or survival (MYH10 and LMNB1), or aberrant tag localization (FBL) (Fig. 1D, 2C-E). One of these clones for each target gene, chosen based on the aggregate result from all quality control assays (Fig. 1G), was chosen for internal imaging studies and public distribution through the Allen Cell Collection at the Coriell Medical Institute Biorepository (https://catalog.coriell.org/1/AllenCellCollection)[47].

**Table 4.** Summary of clones screened and analyzed.

| Experiment | Clones isolated (n) | Clones analyzed (n) | Clones passed all other QC (passed/analyzed) | Final clone chosen for distribution (crRNA used) |
| --- | --- | --- | --- | --- |
| SEC61B | 118 | 58 | 3/3 | cl. 55 (Cr2) |
| FBL | 58 | 20 | 2/2 | cl. 6 (Cr1) |
| ACTB | 192 | 181 | 5/5 | cl. 184 (Cr2) |
| TJP1 | 93 | 91 | 4/4 | cl. 20 (Cr3) |
| PXN | 79 | 68 | 4/5 | cl. 50 (Cr2) |
| LMNB1 | 202 | 93 | 3/3 | cl. 210 (Cr1) |
| TOMM20 | 96 | 83 | 4/4 | cl. 27 (Cr1) |
| DSP | 173 | 164 | 2/3 | cl. 65 (Cr2) |
| TUBA1B | 144 | 138 | 4/5 | cl. 105 (Cr1) |
| MYH10 | 102 | 93 | 5/5 | cl. 80 (Cr4) |
| **Total** | **1257** | **989** | **36/39** |  |

Screen results for all 10 gene tagging experiments. The total number of clones isolated from the FACS-enriched population, clones screened for GFP insertion (analyzed), clones that passed all quality control assays (QC), and the final clone chosen for distribution is shown. The crRNA used to derive the final clone is indicated in parentheses.

### Live cell imaging analyses

Live cell imaging was performed at multiple steps throughout our quality control process starting with the FACS-enriched population of gene edited cells (Fig. 1B, E, F) and then again during the clonal line generation process to ensure proper subcellular localization of the GFP-tagged proteins. Cells were imaged using spinning disk confocal microscopy at low (10×) and high (100×) magnification. Healthy, undifferentiated hiPSCs ranged from 5-20 μm in diameter and 10-20 μm in height and grew in tightly packed colonies (Fig. 3A, B). The observations and subcellular features described below (z-stacks and time lapse movies) can be seen in greater detail in the cell catalog available through the Allen Cell Explorer web portal (http://www.allencell.org/)[48].

**Figure 3.**
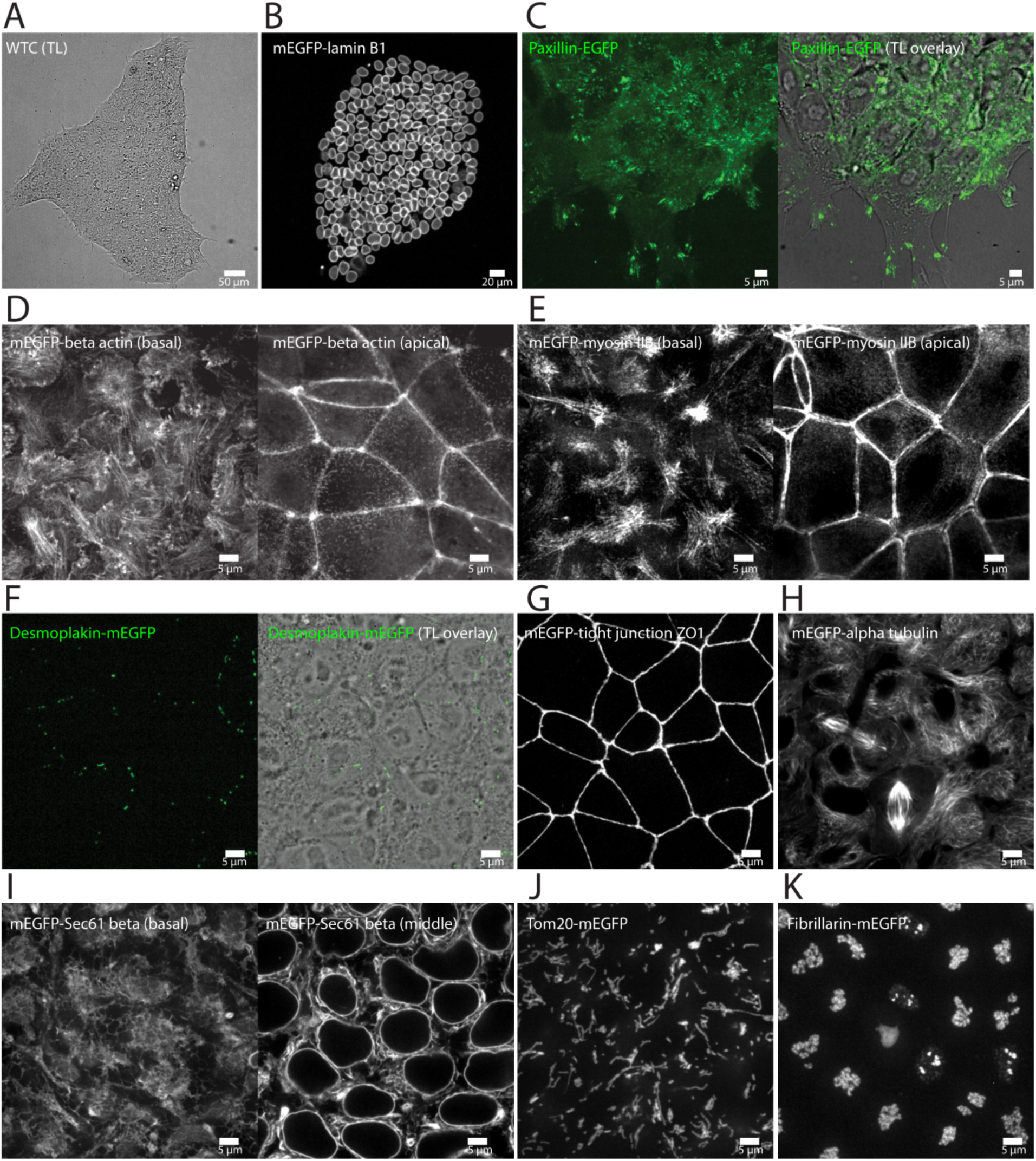
Live cell imaging of final 10 edited clonal lines. Additional images and movies, displaying additional biological features of each cell line, can be found at http://www.allencell.org/. All images are single images taken from a z-stack unless otherwise noted. (**A**) WTC hiPSC stem cell colony (transmitted light image; TL) depicting normal morphology when cells are plated on Matrigel-coated glass. (**B**) mEGFP-tagged nuclear lamin B1 localizes to the nuclear envelope (nuclear periphery) in non-dividing cells and to an extended nuclear lamina within the cytoplasm during mitosis. Image is a maximum intensity projection of the entire colony. (**C**) EGFP-tagged paxillin localizes to puncta at the bottom surface of the cell and larger patches near the dynamic edges of the cell colony, consistent with the localization to matrix adhesions. Some diffuse signal throughout the cytosol is observed. Images are from the bottom of the cells. Right panel is the fluorescence channel overlaid onto the TL channel to indicate colony edges. (**D**) mEGFP-tagged beta actin localizes to stress fibers and lamellipodia at the bottom of the cells (left panel), to a junctional band at the top of cells (right panel), and to regions of cell-cell contact in the center of cells (data not shown). Some diffuse signal throughout the cytosol is consistent with depolymerized actin. (**E**) mEGFP-tagged non-muscle myosin heavy chain IIB localizes basolaterally to stress fibers (left panel), to an apical actin band (right panel) and to regions of cell-cell contact in the centers of cells (data not shown). (**F**) mEGFP-tagged desmoplakin localizes to puncta at apical cell-cell boundaries, consistent with desmosomes. Puncta are not visible in all cells, however when present, there are varying numbers per cell (left panel is a maximum intensity projection of the upper half of the volume of the cells; right panel is the fluorescence channel overlaid onto the TL channel to indicate cell-cell boundaries). (**G**) mEGFP-tagged tight junction protein ZO1 localizes to an apical tight junction band. Weak signal is detectable at cell-cell boundaries in the apicobasal middle of cells (data not shown). Image is a maximum intensity projection. (**H**) mEGFP-tagged alpha tubulin localizes to microtubules, mitotic spindles, primary cilia, and midbodies; some diffuse signal is observed throughout the cytosol, consistent with depolymerized tubulin. (**I**) mEGFP-tagged Sec61 beta is detected in endoplasmic reticulum (ER) sheets and ER tubules throughout the cytoplasm (right panel, image from near the middle of the cells) and in the nuclear periphery (left panel, image from near the bottom of the cells). (**J**) mEGFP-tagged Tom20 localizes to mitochondrial networks throughout the cytoplasm. Image is a maximum intensity projection of 5 z-slices near the bottom of the cells. (**K**) mEGFP-tagged fibrillarin is observed in intra-nuclear structures. Image is a maximum intensity projection. (**A-K**) Scale bars in all panels are as indicated. All imaging was performed in 3D on live cells using spinning disk confocal microscopy with a 100x objective except panels A and B, obtained with a 10x objective. (**B-K**) Representative images of final gene edited cell lines.

We found that several labeled structures exhibited localization patterns reminiscent of the apicobasal polarity in epithelial cells. For example, matrix adhesions (paxillin) formed at substrate contact points with the basal surface of cells and more prominently at the dynamic edges of colonies (Fig. 3C). Filamentous actin (beta actin) was found at the basal surface of colonies both in prominent filaments (stress fibers) and at the periphery of cell protrusions (lamellipodia), as well as in an apical actin band at cell-cell contacts, a feature common in epithelial cells (Fig. 3D). Actomyosin bundles (non-muscle myosin heavy chain IIB) displayed similar basal and apical localization patterns, including at basal stress fibers and an apical band where actin was also observed (Fig. 3D, E). Desmosomes (desmoplakin) localized to distinct puncta at apical cell-cell boundaries, as expected of these junctional complexes in epithelial cells (Fig. 3F). Tight junctions (tight junction protein ZO1) also localized apically to cell-cell junctions (Fig. 3G). These observations suggest the presence of multiple distinct epithelial junction complexes and an overall apical junction zone in edited hiPSC colonies. In addition, microtubules (alpha tubulin) exhibited apicobasal polarity in non-dividing cells, with many microtubules extending parallel to the z-direction as reported for some epithelial cell types (Fig. 3H)[49, 50].

Organelles such as the endoplasmic reticulum (Sec61 beta, Fig. 3I) and mitochondria (Tom20, Fig. 3J) localized throughout the cytoplasm, with greatest density in a cytoplasmic ‘pocket’ near the top of the cell and least density at the center region of the cell that is almost entirely occupied by the nucleus (lamin B1, Fig. 3B). Nucleoli (fibrillarin) localized, as expected, within the nucleus at the apicobasal center of cells (Fig. 3K).

In summary, these observations are consistent with the epithelial nature of tightly packed undifferentiated hiPSCs grown on 2D surfaces. All final candidate clones, spanning 10 editing experiments, exhibited predicted subcellular localization of their tagged proteins (Fig. 3). Taken together, these data demonstrate our ability to identify clonal lines in which genome editing did not interfere with the expected localization of the tagged proteins to their respective structures. Furthermore, live cell time-lapse imaging demonstrated that proper localization occurred throughout cell division and the presence of the tagged protein did not noticeably interfere with cell behavior.

### Cell biological analyses to validate genome editing

To compare the localization of the tagged protein with that of the native, unedited protein, we fixed and immunostained both the edited and unedited cells (Fig. 4A). For all 10 structures, we observed no detectable difference in the appearance of that structure via immunostaining between the unedited and edited line (Fig. 4A, S8). The live, genome-edited cell line generated more interpretable localization data than did fixed cells immunostained for the tagged protein [48]. We also compared endogenously tagged cell lines with unedited cells transiently transfected with constructs expressing the corresponding FP-fusion proteins (GFP or mCherry) and found that the endogenous tag in the edited cell lines greatly enhanced the ability to image these structures as exemplified by mEGFP tagged alpha tubulin, desmoplakin, and Tom20 (Fig. S9). For some proteins, such as alpha tubulin, overexpression via transient transfection led to higher cytoplasmic backgrounds. In endogenously tagged alpha tubulin cells, however, even though the expression levels were lower, the decreased background led to visualization of microtubules with greater clarity (Fig. S9). For other proteins, such as desmoplakin in desmosomes, overexpression led to aberrant localization patterns, such as an increase in the number and size of desmosomes or artificial desmosome-like structures (Fig. S9). At an extreme, overexpression of Tom20 led to poor cell health and loss of normal mitochondrial morphology, while the endogenously tagged cells displayed intact mitochondrial networks with normal morphology and cell viability (Fig. S9). In summary, these genome-edited clonal lines allowed us to observe the tagged proteins and the organelles they represent with exceptional clarity due to endogenous expression and the absence of fixation, staining, and overexpression artifacts.

**Figure 4.**
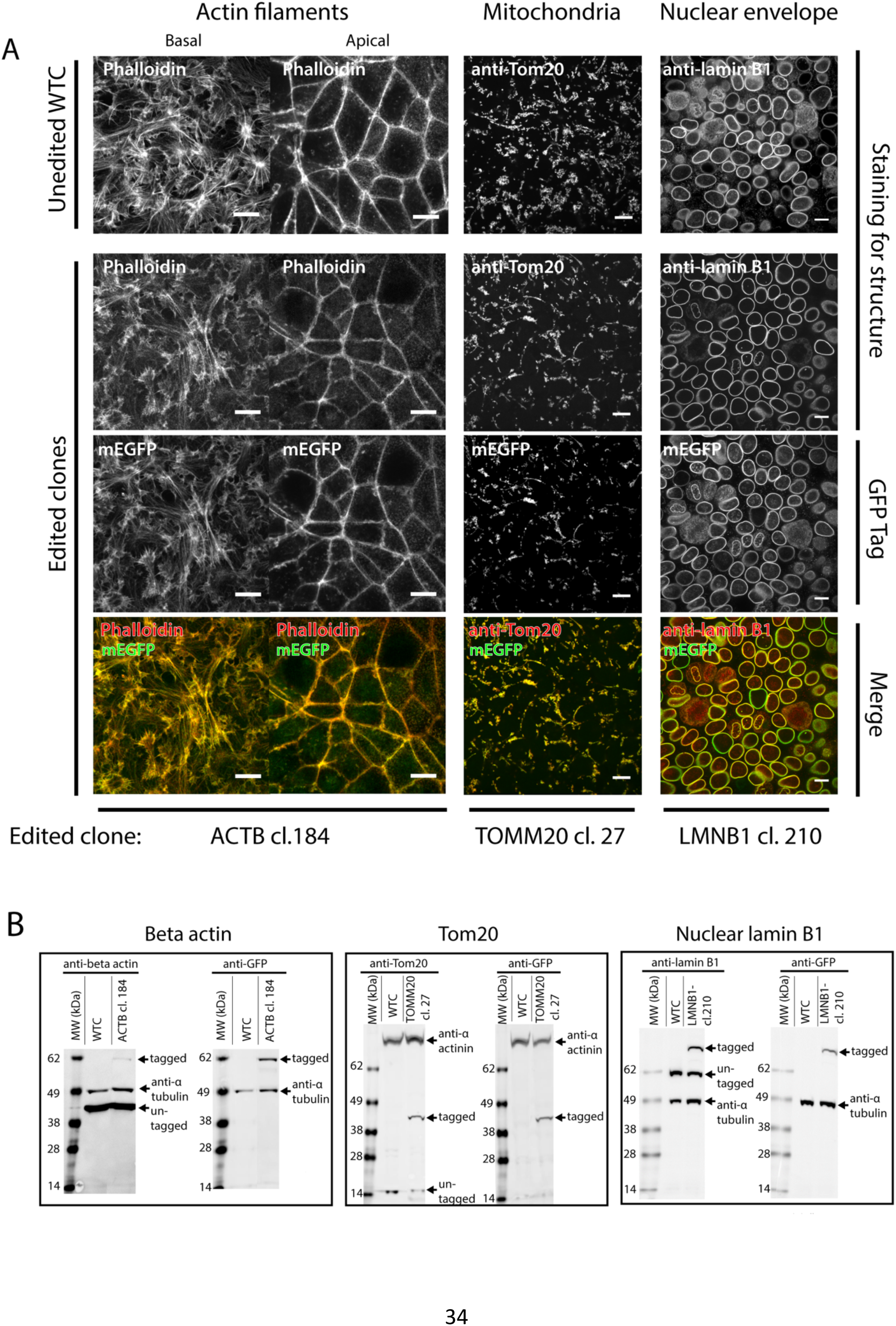
Cell biological assays to evaluate co-expression of tagged and untagged protein forms and their relative contributions to cellular proteome and structure. (**A**) Comparison of labeled structures in edited cells and unedited WTC parental cells. The unedited cells are shown in the top row. Representative images from edited beta actin, Tom20, and lamin B1 are shown as examples (bottom three rows). Cells were stained with Phalloidin-Rhodamine, anti-Tom20 antibody, or anti-lamin B1 antibody, as indicated (Table S4). mEGFP fluorescence (without secondary signal amplification) in genome edited cells and the overlay are also shown (bottom two rows). Scale bars are 10 μm. Additional immunofluorescence data in Fig. S8 and at http://www.allencell.org/. (**B**) Lysate from ACTB cl. 184 (left), TOMM20 cl. 27 (middle), and LMNB1 cl. 210 (right) are compared to unedited WTC cell lysate by Western blot. In all cases, blots with antibodies against the respective proteins (beta actin, Tom20, and lamin B1) are shown in the left blot, and blots with anti-GFP antibodies are shown in the right blot, as indicated. Loading controls were either alpha tubulin or alpha actinin, as indicated.

Western blot analysis was performed on whole cell lysates from candidate edited clones. Immunoblotting with antibodies against the endogenous protein yielded products consistent with both the anticipated molecular weight of the tagged and untagged proteins. The anticipated molecular weight of the tagged protein was further confirmed in all cases by western blot analysis using an anti-GFP antibody (Fig. 4B, S10). Notably, the appropriate GFP fusion protein product was obtained, suggesting that the additional plasmid backbone sequence most likely did not disrupt the coding sequence of the TOMM20 gene, despite our inability to identify a precisely edited TOMM20 clone. Semi-quantitative analysis of the blots was also used to determine the relative abundance of protein products derived from each allele as well as total protein abundance (Table 2). The relative levels of the tagged protein compared to the untagged protein varied by target. While many clones expressed the tagged protein at ∼50% of the total protein in the cell, others did not (Table 2). In the most extreme example, the final tagged beta actin clone expressed only 5% of the tagged protein, but total levels of beta actin in cells were similar to levels found in unedited cells suggesting that the cells were able to compensate for the possible loss of function attributable to tagging while retaining normal viability and behavior.

**Table 2.** Expression analysis of tagged proteins.

| Protein | Clone | Relative level of total protein | Relative level of tagged protein |
| --- | --- | --- | --- |
|  | WTC | 1.00 | N/A |
| Alpha tubulin | cl. 105 | 0.83 | 0.55 |
| Beta actin | cl. 184 | 1.35 | 0.05 |
| Desmoplakin | cl. 65 | n.a | n.a |
| Fibrillarin | cl. 6 | 0.95 | 0.23 |
| Nuclear lamin B1 | cl. 210 | 0.80 | 0.33 |
| Non-muscle myosin heavy chain IIB | cl. 80 | 1.15 | 0.33 |
| Paxillin | cl. 50 | 1.38 | 0.49 |
| Sec61 beta | cl. 55 | 1.47 | 0.53 |
| Tight junction protein ZO1 | cl. 20 | 1.07 | 0.50 |
| Tom20 | cl. 27 | 1.44 | 0.54 |

N/A = not applicable n.a. = not quantified
Relative level of the detected protein in final candidate clones chosen for expansion and distribution compared to unedited WTC cells, is as indicated. The abundance of the tagged protein, relative to the untagged, is as indicated. Separate semi-quantification of untagged and tagged protein versions from the mEGFP-tagged desmoplakin clone was not possible due to the large size of the target protein.

### Stem cell quality control analyses

Upon validating the expression and localization of the GFP-tagged protein in each of the genome-edited lines, we focused on ensuring that each expanded candidate clonal line retained stem cell properties comparable to the unedited WTC cells. Assays included morphology, growth rate, expression of pluripotency markers, and differentiation potential. Undifferentiated stem cell morphology was defined as colonies retaining a smooth, defined edge and growing in an even, homogeneous monolayer (Fig. 5A). Clones with morphology consistent with spontaneous differentiation were rejected [51-54]. Such cultures typically displayed colonies that were loosely packed with irregular edges, with larger and more elongated cells compared to the undifferentiated cells, as observed with one PXN clone (a confirmed bi-allelic edit) (Fig. 5A-right-most image). Although some differences in growth rate were observed in some of the edited clones in preliminary experiments, none were rejected since this was within the variance observed for the unedited cells (data not shown). We also assayed for the expression of established pluripotency stem cell markers including the transcription factors Oct3/4, Sox2 and Nanog, and cell surface markers SSEA-3 and TRA-1-60 (Fig. 5B, Table 3). We found high levels of penetrance in the expression of each marker (>86% of cells) in all final clonal lines from the 10 different genome edits, similar to that of the unedited WTCs (Fig. 5B, Table 3). Consistent with these results, we also found very low penetrance (≤9% of cells) of the early differentiation marker SSEA-1 by flow cytometry in both the edited and control WTC cells (Fig. 5B, Table 3). All 39 clones passed the commonly used guidelines of >85% cells expressing pluripotency markers and <15% cells expressing the differentiation marker SSEA-1 used by various stem cell banks [55].

**Table 3.** Quantitative assessment of pluripotency and cardiomyocyte differentiation for final 10 clonal lines.

| Cell Line | Pluripotency markers, % positive (n) |  |  |  |  |  | Germ layer markers, % positive (n) |  |  | Cardiomyocyte differentiation (n) |  |  |
| --- | --- | --- | --- | --- | --- | --- | --- | --- | --- | --- | --- | --- |
|  | Oct3/4 | Nanog | Sox2 | SSEA-3 | TRA-1-60 | SSEA-1 | Brachyury (mesoderm) | Sox17 (endoderm) | Pax6 (ectoderm) | % cTnT+ | % of expts with beating cells (n) | Range in day of beating initiation |
| Unedited | >95 (7) | >97 (7) | >97 (7) | >97 (7) | >97 (7) | <2 (7) | 84-97 (3) | 47-70 (3) | 95-99 (3) | 70-98 (4) | 89 (38) | d6-d15 |
| PXN cl. 50 | >97 (4) | >99 (4) | >98 (4) | 100 (4) | >86 (4) | <4 (4) | 96 | 58 | 65 | 66-98 (4) | 75 (16) | d7-d17 |
| TUBA1B cl. 105 | >98 (2) | 100 (2) | >99 (2) | 100 (2) | >88 (2) | <6 (2) | 92 | 80 | 89 | 87-96 (4) | 75 (12) | d6-d20 |
| LMNB1 cl. 210 | >95 (2) | >99 (2) | 100 (2) | >98 (2) | >86 (2) | <4 (2) | 93 | 81 | 72 | 39-97 (4) | 80 (10) | d7-d12 |
| TOMM20 cl. 27 | >98 (3) | 100 (3) | >99 (3) | >98 (3) | >91 (3) | <9 (3) | 94 | 82 | 93 | 80-95 (2) | 75 (12) | d7-d13 |
| DSP cl. 65 | >98 (3) | >99 (3) | >99 (3) | >97 (3) | >86 (3) | <5 (3) | 91 | 57 | 78 | 92-95 (2) | 50 (10) | d7-d15 |
| ACTB cl. 184 | 95 | 100 | 100 | 100 | 94 | 8 | >94 (2) | 48 (2) | >98 (2) | 72 | 100 (4) | d7-d8 |
| SEC61B cl. 55 | 97 | 100 | 100 | 95 | 97 | 1 | >97 (2) | 76-79 (2) | >97 (2) | 95 | 80 (5) | d7-d8 |
| FBL cl. 6 | 97 | 100 | 100 | 96 | 91 | 2 | >96 (2) | 57-75 (2) | >98 (2) | 64 | 75 (4) | d7-d9 |
| MYH10 cl. 80 | 96 | 98 | 98 | 100 | 99 | 4 | >94 (2) | 52-64 (2) | 98 (2) | 75-99 (2) | 100 (4) | d7-d16 |
| TJP1 cl. 20 | 100 | 99 | 100 | 100 | 94 | 4 | >94 (2) | 53-66 (2) | >99 (2) | 75-93 (3) | 100 (6) | d7-d17 |

Final clones chosen for each gene distribution were assessed for pluripotency, embryonic germ layer, and cardiomyocyte differentiation potential by flow cytometry. The percentage of stained cells above the positive staining threshold for the indicated marker is listed with the number of trials shown as (n) where more than one trial was performed. Values reflect either the lowest measured percentage (when indicated by >), the highest measured percentage (when indicated by <), or the range observed across multiple experiments. Unedited WTC samples accompanied edited cell lines in all trials. The fraction of cTnT-positive cells reported is from trials with confirmed contractile cells.

To confirm the pluripotency of the clones, we compared the potential for germ layer differentiation between unedited cells and one edited clonal line for each of the 10 targeted structures. Each cell line was differentiated for 5-7 days under defined conditions to mesoderm, endoderm, and ectoderm using lineage-specific media for each lineage. The cells were stained for early markers of germ layer differentiation (Brachyury, Sox17, and Pax6) and analyzed by flow cytometry (Fig. 5C, Table 3) [56-59]. While the differentiation into each germ layer was variable between clones, all three germlayer markers in the edited clones showed increased expression relative to undifferentiated cells (Fig. 5C). In all edited clones tested, ≥91% of cells expressed Brachyury after mesodermal differentiation, ≥48% expressed Sox17 after differentiation to endoderm, and ≥65% expressed Pax6 upon ectoderm differentiation (Fig. 5C, Table 3).

As a third measure of pluripotency, we evaluated differentiation into a more developmentally advanced cellular fate by interrogating each clone’s ability to differentiate into cardiomyocytes. We followed an established differentiation protocol using a combination of growth factors and small molecules and evaluated cultures for spontaneous beating (days 6-20) and cardiac Troponin T (cTnT) expression (days 20-25), an established marker for cardiac fate [60, 61]. Due to heterogeneity in these differentiation assays, a clone was only validated for differentiation if spontaneous beating was observed in at least 3 experiments (the total number of experiments reported reflect the status of our gene editing and differentiation pipeline at the time of manuscript preparation). All 10 clonal lines displayed successful cardiomyocyte differentiation with spontaneous beating and cTnT expression comparable with the parental WTC line (Fig. 5D, E, Table 3). During our quality control process, we tested a total of 39 edited lines of which 38 displayed successful differentiation. The single exception was one of three tested DSP clones which did not meet our criterion for spontaneous beating. While the other two clones from this experiment passed our criteria, they nevertheless showed reduced propensity for beating (50% experiments with beating for clone 65, Table 3) compared to the other edited clones. Since desmoplakin is known to be an important structural protein in cardiomyocytes, this may indicate a sensitivity to tagging that does not emerge until differentiation [62, 63]. In general, the pluripotency, germ layer differentiation potential, and cardiomyocyte differentiation data support the conclusion that fusing GFP with these endogenously expressed proteins does not typically disrupt pluripotency.

In summary, of the 39 clones analyzed for genomic stability, expression of the tagged protein, proper subcellular localization, stem cell morphology, and pluripotency, only 3 clones were rejected due to changes in either stem cell morphology (only the bi-allelically edited PXN clone), abnormal karyotype (1 of 5 TUBA1B clones), and impaired differentiation into cardiomyocytes (1 of 3 DSP clones) (Table 4). Therefore, these data underscore the ability of hiPSCs to not only tolerate mono-allelic GFP tags in key proteins, but also retain properties of highly pluripotent stem cells. Of the 36 fully validated clones, 20 (2 per gene/structure) were expanded for banking and re-analyzed for genomic stability by karyotype analysis and sterility. Final clonal lines encompassing all 10 structures described in this report are now openly available to the research community [47].

## Discussion

While several studies demonstrate that genome editing in stem cells is feasible using CRISPR/Cas9, the reliability and efficiency of HDR-mediated “knock-in” delivery of large sequences, such as GFP, at diverse genomic loci is less clear. This is due to the scarcity of studies in which multiple loci are edited within a given biological context. Using hiPSCs and the methods described here, we have successfully edited 10 diverse target genes. This approach and accompanying analyses may provide insight for editing novel targets. We have generated 36 edited clonal lines that satisfy our cell biological and stem cell quality criteria from 10 different genome editing experiments. The finding that >90% (36/39) of our final GFP-tagged lines meet our stringent genomic and cell biological quality control criteria suggests that genome editing in stem cells is feasible and that these edited cells often tolerate a tagged endogenous protein.

The systematic nature of our study, which included a standardized editing strategy and a quality control workflow, the diversity of genomic loci targeted, and the large number of clones screened led to several key observations that are summarized below: First, genes with sufficient expression in stem cells to permit successful enrichment (by FACS) can be effectively edited with this methodology. This may also include genes with relatively modest expression and/or punctate localization as seen for PXN and DSP. This approach may therefore be useful for tagging a broad range of cellular proteins.

Second, high editing efficiency is not required for successful tagging as observed for many of the genes described here. Experiments in which HDR was inefficient, such as MYH10 (0.1%), nevertheless yielded precisely edited clones that satisfied both genomic and cell biological quality control criteria. Considerable discussion has been devoted to the inefficiency of HDR in human cells in general, and stem cells in particular, and whether this limitation can be ameliorated [26, 64, 65]. Our data suggests that successful editing is feasible regardless of the observed editing rate, and that the true limitation is the ability to enrich for the edited cells.

Third, testing multiple crRNAs led to a high likelihood of successful editing and subsequent identification of precisely edited clones. We did not observe any predictable trends with regard to achieving greater overall editing success among crRNAs based on features such as the proximity of their PAM (-3 position) to the desired point of insertion. Despite our preference for clones independently derived from multiple crRNAs so that off-target concerns are mitigated, this was only possible in 2 out of 10 experiments (DSP and SEC61B). In all other experiments, all final validated clones were derived from one crRNA with the other crRNAs failing due to absence of or imprecise editing, mislocalization of signal, or altered stem cell morphology. Since experiments with multiple crRNAs were performed in parallel, we attribute many of these differences to locus and crRNA sequence-specific effects on HDR efficiency and precision. Therefore, testing several crRNAs helped us achieve successful HDR with at least one crRNA for all 10 gene targets and we recommend this strategy.

Fourth, prioritizing crRNA design according to anticipated off-target sequences may mitigate undesired off-target editing. Although we did not perform unbiased genome-wide off target analysis, we did not observe any mutations in 39 edited clones at 142 high homology sites. This may be due in part to our rejection of crRNAs with predicted promiscuous off-target profiles. Since our data also suggest that a preferred crRNA cannot necessarily be inferred at the outset of an experiment, as discussed above, we recommend that several crRNAs with the most favorable predicted off-target profile be chosen for editing. We also minimized the impact of potential unknown off-target events by screening multiple clonal lines across a broad range of genomic and cell biological criteria.

Fifth, the relatively modest rate of NHEJ observed in untagged alleles (Fig. 2E-77% wild type) with an exact on-target crRNA binding sequence and our inability to detect off-target mutations at candidate off-target sites suggest that off-target NHEJ events are likely minimal with this editing approach. We hypothesize that use of an RNP protocol with relatively low concentrations of the editing components for “gentler” editing may restrict both on- and off-target mutagenesis via NHEJ. We are aware of several investigations using plasmid-based CRISPR/Cas9 overexpression platforms where the observed NHEJ rate is in excess of 50% at multiple loci [66] (personal communication, Bruce Conklin and Jacob Corn). The hypothesis that our editing method results in modest genomic insult is additionally supported by the bias toward mono-allelic edits generated with our methods. Alternatively, it is possible that we obtained almost exclusively mono-allelic edits due to lethality associated with bi-allelic tagging since our target genes encode proteins representing key cellular structures such as the actin and microtubule cytoskeleton, mitochondria, and nuclear envelope. Although bi-allelic editing was not a high priority for our purposes of live cell imaging, increasing the effective “dose” of the various components of the CRISPR/Cas9 system may represent a route to achieving greater rates of bi-allelic editing, provided the cells can tolerate two modified alleles.

Sixth, genetic validation is crucial prior to using edited cells in downstream applications. Clones with integrated plasmid backbone were prevalent and in some cases would have been overlooked if not for careful screening in our ddPCR assay. Therefore, we strongly advocate for thorough genetic screening of edited cells that includes direct detection of the plasmid donor backbone in the host cell genome. In addition, several edited populations (TJP1 Cr1, ACTB Cr1, TUBA1 Cr2) and clones (TOMM20), while incorporating the GFP at the correct locus did so imprecisely, resulting in relatively large insertions most likely in the non-coding regions flanking the GFP such as the 3′UTR (TOMM20). These edited cells displayed GFP localization to the correct subcellular structure, suggesting that imprecise editing did not affect the coding region of the targeted gene (e.g.TOMM20). However, this does not rule out the possibility of disrupting potential regulatory elements, if present, at the site of the plasmid backbone insertion. Our data suggests that these insertions may in fact commonly represent complete duplications of the donor plasmid cassette, which may confound junctional PCR analysis, underscoring the potential impact of these deceptive HDR outcomes in GFP tagging experiments.

Finally, the fundamental functions of the proteins tagged in our collection of edited lines and the broad potential applications we envision for them required quality control assays to ensure the endogenous GFP tag has no obvious adverse effects. This was crucial for all edited clones, especially when imprecise editing was detected and mis-regulation of the target gene was a greater concern. To address some of these concerns, we performed several quality control assays intended to identify whether the GFP tag disrupted cellular processes. These included imaging of live cells to confirm localization of the tagged protein to the appropriate structure and comparison of structures labeled in fixed edited and unedited cells. These imaging studies indicate the GFP tag was well tolerated. Western blot analysis of all 10 fusion proteins confirmed expression of the predicted protein and normal processing in all cases, including Tom20, where plasmid sequence was incorporated into the endogenous locus. These experiments also provided an indication of the relative and total abundance of both the tagged and untagged protein forms, which varied significantly between experiments. In one notable case, ∼95% of the beta actin in the cell was the untagged form with no decrease in the total amount of beta actin, relative to unedited cells. This illustrated the ability of hiPSCs to differentially regulate the expression of the tagged and untagged proteins. In cases such as beta actin, which is known for its difficulty in accommodating a large tag in non-mammalian systems, this ability is noteworthy and useful [67, 68]. Karyotyping and pluripotency assays further confirmed that these edited cells retained both genomic stability and differentiation potential. Therefore, we conclude that WTC hiPSCs, and potentially other cells, are encouragingly tolerant of mono-allelic gene tagging at diverse loci to the extent that we have characterized them.

In conclusion, we produced edited hiPSC clones spanning 10 target genes that met the above-described criteria. These clones were comparable to unedited cells despite harboring endogenous gene tags in diverse proteins. We also observed that clones with precise editing showed GFP localization of the fusion protein to the appropriate organelle with remarkable consistency. We are encouraged by the high tolerance for mono-allelic tagging and the absence of significant changes in the observed biology of clones expressing the edited genes. Live imaging of the resulting 10 edited lines clearly demonstrated key features of the tagged structure in unprecedented image clarity. Studies utilizing these lines will likely benefit from both the absence of overexpression and staining artifacts, and the use of healthy, diploid cells to reveal novel aspects of cell biology [69]. Since this is the first time many of these structural genes have been endogenously tagged in human stem cells, the quality control assays we have conducted instill confidence in these observations and the resulting insights into the organization and activities of hiPSCs. We have made these cell lines and plasmids used to derive them available to the research community through the Allen Cell Collection [47] and the Allen Cell Collection Plasmids [70].

## Materials and Methods

### Cell Culture

All work with human induced pluripotent stem cell (hiPSC) lines was approved by internal oversight committees and performed in accordance with applicable NIH, NAS, and ISCCR Guidelines. The WTC hiPSC cell line was generated by Bruce Conklin [21]. Upon receipt, we authenticated the cell line with the donor fibroblasts using short tandem repeat (STR) analysis (Cell Line Genetics). Edited cell lines described in this report can be obtained by visiting the Allen Institute for Cell Science website [48]. WTC hiPSCs were cultured in a feeder-free system on tissue culture dishes or plates coated with GFR Matrigel (Corning) diluted 1:30 in cold DMEM/F12 (Gibco). Undifferentiated cells were maintained with mTeSR1 media (STEMCELL™ Technologies, Inc.) supplemented with 1% (v/v) Penicillin-streptomycin (P/S) (Gibco). Cells were not allowed to reach confluency greater than 85% and were passaged every 3-4 days by dissociation into single-cell suspension using StemPro^®^ Accutase (Gibco). When in single cell suspension, cells were counted using a Vi-CELL-XR^®^ Series Cell Viability Analyzer (Beckman Coulter). After passaging, cells were re-plated in mTeSR1 supplemented with 1% P/S and 10 μM ROCK inhibitor (Stemolecule Y-27632, Stemgent) for 24 h. Media was replenished with fresh mTeSR1 media supplemented with 1% P/S daily. Cells were maintained at 37°C and 5% CO_2_. A detailed protocol can be found at the Allen Institute for Cell Science web portal [48].

### Donor Plasmids, crRNAs and Cas9 protein

Donor plasmids were designed uniquely for each target locus, with each following a similar design strategy. Homology arms 5′ and 3′ of the desired insertion site were each 1 kb in length and designed from the GRCh38 reference genome. WTC-specific variants (SNPs and INDELs) were identified from publicly available exome data [22]. In cases where the WTC-specific variant was heterozygous, the reference genome variant was used in the donor plasmid; when the WTC-specific variant was homozygous, the WTC-specific variant was used in the donor plasmid. Linkers for each protein were unique and were used to join the terminus of the protein with the GFP-encoding sequence (inserted 5′ of GFP for C-terminal tags and 3′ of GFP for N-terminal tags). To prevent crRNAs from targeting donor plasmid sequence, mutations were introduced to disrupt Cas9 recognition or crRNA binding; when possible, these changes did not affect the amino acid sequence. All plasmids and design criteria used to construct them are available at Addgene (https://www.addgene.org/The_Allen_Institute_for_Cell_Science)[70]. The plasmids were initially created either by In-Fusion (Clontech) assembly of gBlock pieces (IDT) into a pUC19 backbone (New England Biolabs), or were synthesized and cloned into a pUC57 backbone by Genewiz. Plasmid DNA for transfection was prepared using endotoxin-free purification kits (NucleoBond^®^ Xtra Maxi EF, Clontech). Custom synthetic crRNAs and their corresponding tracrRNAs were ordered from either IDT or Dharmacon. Recombinant wild type *S. pyogenes* Cas9 protein was purchased from the UC Berkeley QB3 Macrolab. All tagging experiments discussed in the current report used the mEGFP (K206A) sequence, except PXN, which was performed with EGFP instead. Detailed information on editing design can be found at The Allen Cell Explorer [48].

### Transfection and Enrichment by Fluorescence-Activated Cell Sorting (FACS)

Cells were dissociated into single-cell suspension using Accutase as previously described. Transfections were performed using the Neon transfection system (Life Technologies). We evaluated various delivery methods including CRISPRMax (ThermoFisher), GeneJuice^®^ transfection reagent (EMD Millipore), Amaxa (Lonza) and Neon (ThermoFisher) and concluded that Neon electroporation resulted in favorable co-introduction of protein, RNA, and plasmid into hiPSCs as measured by transfection of a control reporter plasmid and T7 assays as a readout for Cas9 activity (data not shown). Cas9:crRNA:tracrRNA precomplexed 1:1:1 and cotransfected with 2 μg of donor plasmid optimally balanced editing efficiency with cell survival after transfection (data not shown) and we chose this platform for all editing experiments. A detailed protocol for the RNP transfection can be also be found at the Allen Cell Explorer web portal [48].

A cell pellet of 8×10^5^ cells was resuspended in 100 μL Neon Buffer R with 2 μg donor plasmid, 2 μg Cas9 protein, and duplexed crRNA:tracrRNA in a 1:1 molar ratio to Cas9. Prior to addition to the cell suspension, the Cas9/crRNA:tracrRNA RNP was precomplexed for a minimum of 10 minutes at room temperature. Electroporation was with one pulse at 1300 V for 30 ms. Cells were then immediately plated onto Matrigel-coated 6-well dishes with mTeSR1 media supplemented with 1% P/S and 10 μM ROCK inhibitor. Transfected cells were cultured as previously described for 3-4 days until the transfected culture had recovered to ∼70% confluence.

Cells were harvested for FACS using Accutase as previously described. The cell suspension (0.5 – 1.0 × 10^6^ cells/mL in mTeSR1 with ROCK inhibitor) was filtered through a 35 μm mesh filter into polystyrene round bottomed tubes. Cells were sorted using a FACSAriaIII Fusion (BD Biosciences) with a 130 μm nozzle and FACSDiva software (BD Biosciences). Forward scatter and side scatter (height versus width) were used to exclude doublets and the GFP-positive gate was set using live, untransfected WTC cells such that <0.1% of untransfected cells fell within the gate. Sorted populations were plated into Matrigel-coated 96-well plates (<2 × 10^3^ cells recovered) or 24-well plates (<1×10^4^ cells recovered) for expansion of the whole enriched population before clone isolation. In some cases, (e.g. PXN), co-isolation of presumptively unedited cells was tolerated due to the weak GFP fluorescence intensity of tagged protein. To determine %HDR, data were analyzed using FlowJo V.10.2 (TreeStar, Inc.).

### Clonal Cell Line Generation

FACS-enriched populations of edited cells were seeded at a density of 1×10^4^ cells in a 10 cm Matrigel-coated tissue culture plate. After 5-7 days clones were manually picked with a pipette and transferred into individual wells of 96-well Matrigel-coated tissue culture plates with mTeSR1 supplemented with 1% P/S and 10 μM ROCK inhibitor for 1 day. After 3-4 days of normal maintenance with mTeSR1 supplemented with 1% P/S, colonies were dispersed with Accutase and transferred into a fresh Matrigel-coated 96-well plate. After recovery, the plate was divided into daughter plates for ongoing culture, freezing, and gDNA isolation.

To cryopreserve clones in a 96-well format, when cells were 60-85% confluent they were dissociated and pelleted in 96-well V-bottom plates. Cells were then resuspended in 60 μL mTeSR1 supplemented with 1% P/S and 10 μM ROCK inhibitor. Two sister plates were frozen using 30 μL cell suspension per plate, added to 170 μL CryoStor^®^ CS10 (Sigma) in non-Matrigel coated 96-well tissue culture plates. Plates were sealed with Parafilm and stored at - 80°C.

### Genetic Screening with droplet digital PCR (ddPCR)

During clone expansion, >1500 cells were pelleted from a 96-well plate for total gDNA extraction using the PureLink Pro 96 Genomic DNA Purification Kit (Life Technologies). ddPCR was performed using the Bio-Rad QX200 Droplet Reader, Droplet Generator, and QuantaSoft software. The reference assay for the 2-copy, autosomal gene RPP30 was purchased from Bio-Rad. For primary ddPCR screening the assay consisted of three hydrolysis probe based PCR amplifications targeted to three different genes: GFP (insert), AMP or KAN (backbone), and the genomic reference RPP30. The following primers were used for the detection of GFP (5′- GCCGACAAGCAGAAGAACG-3′, 5′-GGGTGTTCTGCTGGTAGTGG-3′) and hydrolysis probe (/56-FAM/AGATCCGCC/ZEN/ACAACATCGAGG/3IABkFQ/). This assay was run in duplex with the genomic reference RPP30-HEX. The PCR for detection of the AMP gene used the primers (5′- TTTCCGTGTCGCCCTTATTCC -3′, 5′- ATGTAACCCACTCGTGCACCC -3′) and hydrolysis probe (/5HEX/TGGGTGAGC/ZEN/AAAAACAGGAAGGC/3IABkFQ/).

The PCR for detection of the KAN gene used the primers (5′-AACAGGAATCGAATGCAACCG-3′, 5′-TTACTCACCACTGCGATCCC-3′) and hydrolysis probe ((/5HEX/GTGAAAATA/ZEN/TTGTTGATGCGCTGG/3IABkFQ/).

PCR reactions were prepared using the required 2x Supermix for probes with no UTP (Bio-Rad) with a final concentration of 400 nM for primers and 200 nM for probes, together with 10 units of HindIII and 3 μL of sample (30-90 ng DNA) to a final volume of 25 μL. Each reaction prior to cycling was loaded into a sample well of an 8-well disposable droplet generation cartridge followed by 70 μL of droplet generator oil into the oil well (Bio-Rad). Droplets were then generated using the QX200 droplet generator. The resulting emulsions were then transferred to a 96 well plate, sealed with a pierceable foil seal (Bio-Rad), and run to completion on a Bio-Rad C1000 Touch thermocycler with a Deep Well cycling block. The cycling conditions were: 98°C for 10 min, followed by 40 cycles (98°C for 30 s, 60°C for 20 s, 72°C for 15 s) with a final extension at 98°C for 10 min. After PCR, droplets were analyzed on the QX200 and data analysis was preformed using QuantaSoft software.

The reported final copy number of GFP per genome was calculated as the ratio of [(copies/μL_GFP_)-(copies/μL_nonintegratedAMP_)]/(copies/μL_RPP30_), where a ratio of 0.5 is equal to 1 integrated copy/genome and a ratio of 1 is equal to 2 integrated copies/genome. The AMP or KAN signal was determined to be from residual non-integrated/background plasmid when the ratio of AMP/RPP30 or KAN/RPP30 fell below 0.2 copies/genome, as ratios above 0.2 copies/genome could indicate a non-clonal population with at least 1 copy of the backbone stably integrated. For primary screening the ratios of (copies/μL_GFP_)/(copies/μL_RPP30_) were plotted against [(copies/μL_AMP_)/(copies/μL_RPP30_) to identify cohorts of clones for ongoing analysis. A detailed protocol for these methods is available at the Allen Cell Explorer web portal [48].

### Genetic Screening with tiled junctional PCR

PCR was used to amplify the tagged allele in two tiled reactions spanning the left and right homology arms, the GFP and linker sequence, and portions of the distal genomic region 5′ of the left homology arm and 3′ of the right homology arm (Fig. 2) using gene-specific primers (Table S3). Both tiled junctional PCR products were Sanger sequenced (Genewiz) bidirectionally with PCR primers when their size was validated by gel electrophoresis and/or fragment analysis (Advanced Analytics Technologies, Inc. Fragment Analyzer). In final clones, a single, non-tiled junctional PCR reaction using the gene-specific external 5′ and 3′ junctional primers (Table S3) was used to amplify both the edited and wild type allele in a single reaction. All PCR reactions described above were prepared using PrimeStar® (Takara) 2x GC buffer, 200 μM DNTPs, 1 unit PrimeStar HS polymerase, 800 nM primers, 10 ng gDNA in a final volume of 25 μL. Cycling conditions were as follows (98°C 10s, 70°C 5s, 72°C 60s) x 6 cycles at -2°C/cycle annealing temperature, (98°C 10s, 54°C 5s, 72°C 60s) × 32 cycles, 12°C hold.

### Screening for clones with wild type untagged allele sequences

PCR was also used to amplify the untagged allele using gene-specific primers (Table S3). These primers did not selectively amplify the unmodified locus, as was the case for tiled junctional PCR amplification of the tagged allele, but rather amplified both untagged and tagged alleles. PCR was performed with the same Primestar reagents and cycling conditions as described above. Tracking of insertions and deletions (INDELs) by decomposition (TIDE) analysis was performed manually on the amplification reaction after bidirectional Sanger sequencing in order to determine the sequence of the untagged allele. For all final clones with wild type untagged alleles, the PCR product corresponding to the untagged allele was gel isolated and sequenced to confirm the initial result from TIDE analysis.

### Off-target PCR screening

Cas-OFFinder was used to identify potential off-targets (NRG PAMs with up to 3 mismatches and 1 DNA or RNA bulge) in GRCh38 genome build (GCA_000001405.15). Cas-OFFinder output was further filtered to identify the most problematic off-targets with the fewest number of flaws (flaw = mismatch or bulge) relative to the on-target crRNA. Problematic off-targets were defined as off-targets with up to one flaw in the seed region (10 nts at 3′ end) and up to 2 flaws in the non-seed region (10 nts at 5′ end) with an NGG or NAG PAM (Fig. S2) [40]. 8-10 of these off-targets were selected for sequencing with the goal of checking ∼4 off-targets that fell close to exons (within 50 bp) or within exons (exon feature in GRCh38 NCBI annotation 107) and ∼4 off-targets that were closest in sequence to on-target crRNA. Approximately 300 bp of sequence flanking each off-target was amplified by PCR and Sanger sequenced.

### Off-target categorization

We categorized off-targets based on their PAM -3 location (expected double strand break site): exonic =PAM-3 site (expected double strand break site) inside exon or within 50 bp of exon; genic = not in exonic category but either inside gene (intron) or within 200 bp of gene; non-genic = outside of exons or genes). The locations of exons and genes were identified from GRCh38 assembly (GCA_000001405.15) NCBI annotations version 107. We also categorized off-targets based on their sequence profile: number of flaws (mismatches or DNA/RNA bulges), location of flaws (seed vs non-seed), and PAM sequence (NGG vs NAG) (Fig. S2). The same off-target site (same = same strand and same double strand break site) can have a slightly different sequence depending on the position of the bulge and the ability to accommodate either a bulge or a mismatch flaw, and Cas-OFFinder will report this same site multiple times. We collapsed these multiple outputs into a single off-target site and used the sequence profile ranking scheme in Fig. S2 to categorize the resulting off-target.

### Cell Plating for Imaging

Cells were plated on glass bottom multi-well plates (1.5H glass, Cellvis) coated with phenol red-free GFR Matrigel (Corning) diluted 1:30 in phenol red-free DMEM/F12 (Gibco). Cells were seeded at a density of 2.5x10^3^ in 96-well plates and 12.5×10^3^ to 18×10^3^ on 24-well plates and fixed or imaged 3-4 days later. A detailed protocol can be found at the Allen Cell Explorer web portal [48].

### Live Cell Imaging

Cells were maintained with phenol red free mTeSR1 media (STEMCELL Technologies) one day prior to live cell imaging. Cells were imaged on a Carl Zeiss spinning disk microscope with a Carl Zeiss 20×/0.8 NA plan APOCHROMAT or 100×/1.25 W C-APOCHROMAT Korr UV Vis IR objective, a CSU-X1 Yokogawa spinning disk head, and Hammamatsu Orca Flash 4.0 camera. Microscopes were outfitted with a humidified environmental chamber to maintain cells at 37°C with 5% CO_2_ during imaging.

### Immunocytochemistry and Fixed Cell Imaging

All cell lines except for TUBA1B were fixed and permeabilized in 24- or 96-well plates with a solution of 4% paraformaldehyde (Electron Microscopy Sciences Hatfield, PA) and 0.5% Triton X-100 (EMD Millipore) for 10-15 min. TUBA1B cells were fixed in -20°C methanol for 5 min. Following fixation and permeabilization, all cells were blocked with 1% or 5% BSA (ThermoFisher Scientific) in 1× PermWash Buffer (BD Biosciences), incubated in primary antibody overnight at 4°C, followed by incubation in AlexaFluor^®^-conjugated secondary antibodies (Table S4) and DAPI (1× NucBlue Fixed Cell Stain, ThermoFisher Scientific) for 2hr at room temperature. In the case of ACTB, cells were stained with Rhodamine Phalloidin (ThermoFisher Scientific) at 1:1000. SlowFade Gold (ThermoFisher Scientific) mounting media was added to the cells after a final washing step and cells were stored at -20°C until imaged as described in the above section. All fixed cells except for DSP and SEC61B were imaged on a 3i system with a Zeiss 100×/1.46 NA alpha-plan APOCHROMAT oil objective, CSU-W1 Yokogawa spinning disk head and Hammamatsu Orca Flash 4.0 camera. DSP cells (and corresponding controls) were imaged on a Zeiss laser scanning (LSM) 880 confocal microscope with a Zeiss C-APOCHROMAT 40×/1.2 W Korr FCS M27 objective. SEC61B cells were imaged with a Zeiss 100×/1.25 W C APOCHROMAT Korr UV Vis IR objective, a CSU-W1 Yokogawa spinning disk head, and Hammamatsu Orca Flash 4.0 camera.

### Western Blotting

Whole-cell lysate was extracted from cell lines using M-PER buffer (ThermoFisher Scientific) supplemented with 1X Halt protease and phosphatase inhibitors (ThermoFisher), 0.16 M MgCl_2_ and 100 U Pierce universal nuclease (ThermoFisher Scientific) on ice for 30 min or urea sample buffer (USB: 8M urea, 1% SDS, 10% glycerol, 0.06 M Tris pH 6.8, 0.001% pyronin Y). M-PER-based cell lysates were boiled for 5 minutes, then diluted with 1:1 Bolt LDS running buffer (ThermoFisher Scientific), heated at 70°C for 10 minutes and stored at -20°C. Prior to gel electrophoresis, M-PER-based samples were diluted to a final concentration with 1X Bolt LDS running buffer, supplemented with 1X Bolt sample reducing agent (Thermo Fisher Scientific) and heated to 70°C for 10 minutes. USB-based cell lysates were triturated with a syringe and a 27g needle and stored at -20°C. For electrophoresis, USB-based samples were supplemented with 5% beta mercaptoethanol. Lysates were separated on 4-12% Bolt Bis-Tris Plus gels (ThermoFisher Scientific), 1X MOPS LDS running buffer (ThermoFisher Scientific) supplemented with Bolt antioxidant (ThermoFisher Scientific) or NuPAGE Novex 3-8% Tris-Acetate gels (ThermoFisher Scientific), 1× Tris acetate SDS running buffer (ThermoFisher Scientific), supplemented with Bolt antioxidant and transferred onto 0.45 μm nitrocellulose membranes (ThermoFisher Scientific) using the XCell SureLock Mini-Cell System (ThermoFisher Scientific). Untagged and GFP-tagged proteins were detected using protein-specific and GFP-specific antibodies in PBS-T with 5% milk overnight at 4°C (see Table S4 for details). Beta actin was used as the loading control for PXN, SEC61B, TUBA1B, TJP1, and MYH10, while alpha tubulin was used for LMNB1, FBL, ACTB and DSP, and alpha actinin for TOMM20 (Table S4). Goat polyclonal anti-mouse AlexaFluor^R^ 647-conjugated secondary antibody or goat polyclonal anti-rabbit AlexaFluor^R^ 647-conjugated antibody) were used as secondary antibodies as described in Table S4. Blots were imaged at different exposure times using the ChemiDoc MP system (Bio-Rad Laboratories) and appropriate exposure times were used for semi-quantitative analysis of protein levels.

### Transient Transfection for Live Cell Imaging

The constructs pmEGFP-a-tubulin-C1 (gift from Daniel Gerlich, addgene plasmid # 21039), 1136-Desmoplakin-GFP (gift from Kathleen Green, addgene plasmid # 32227) and mCherry-TOMM20-N-10 (gift from Michael Davidson, addgene plasmid #55146) were used for transient transfections [29, 71]. Bacterial strains containing the constructs were grown in 200 mL Terrific Broth (ThermoFisher Scientific) supplemented with 100 μg/mL carbenicillin (ThermoFisher Scientific) or 100 μg/mL kanamycin (ThermoFisher Scientific) overnight in a shaking incubator at 37°C. DNA plasmids were extracted from the bacterial cultures using an EndoFree Plasmid Maxi Kit (QIAGEN) according to manufacturer’s instructions and resuspended in the provided endotoxin-free water. DNA concentrations were quantified using a NanoDrop8000 (ThermoFisher Scientific) and stored at - 20°C. A working DNA stock for transfections was made by diluting the DNA to a final concentration of 0.25 μg/μL in Opti-MEM (ThermoFisher Scientific) and stored at -20°C. Three days after cells were plated, they were transfected using the GeneJuice^®^ Transfection Reagent (adjusted as discussed above) (EMD Millipore). Media was replaced with phenol red-free mTeSR1 30-60 minutes prior to transfection. 1.5 μL GeneJuice transfection reagent was diluted in 25 μL Opti-MEM (ThermoFisher Scientific) and incubated at room temperature for 5 minutes. 1 μg DNA was added to the GeneJuice-Opti-MEM solution and incubated for 10 minutes. 6 μL of this final transfection solution was added per 100 μL media in the well of a 96 well-plate. Live cells were imaged as specified above one day after transfection.

### Pluripotency Testing by Flow Cytometry

Cells were dissociated with Accutase as previously described, fixed with CytoFix Fixation Buffer™ (BD Biosciences) for 30 minutes at 4°C, and frozen at -80°C in KnockOut™ Serum Replacement (Gibco) with 10% DMSO. After thawing, cells were washed with 2% BSA in DPBS and half of the cells were stained with anti-TRA-1-60 Brilliant Violet™ 510, anti-SSEA-3 AlexaFluor^®^ 647, and anti-SSEA-1 Brilliant Violet™ 421 (all BD Biosciences) for 30 minutes at room temperature, then washed with 2% BSA in DPBS. The other half of the cells were permeabilized with 0.1% Triton X-100 and 2% BSA in DPBS for 30 minutes at room temperature followed by staining with anti-Nanog AlexaFluor^®^ 647, anti-Sox2 V450, and anti-Oct3/4 Brilliant Violet™ 510 (all BD Biosciences) for 30 minutes at room temperature. Intracellularly stained cells were washed with BD Perm/Wash™ buffer (BD Biosciences). All cells received a final wash of 2% BSA in DPBS and before being resuspended in 2% BSA in DPBS for acquisition. Cells were acquired on a FACSAriaIII Fusion (BD Biosciences) equipped with 405, 488, 561, and 637 nm lasers and analyzed using FlowJo V.10.2 (Treestar, Inc.). Approximately one fourth of samples were stained with a panel designed for a different FACSAriaIII configuration: anti-TRA-1-60 PerCP-Cy™5.5, anti-SSEA-3 PE, anti-SSEA-1 Brilliant Violet™ 650, anti-Nanog PE, anti-Sox2 AlexaFluor^®^ 647 and anti-Oct3/4 PerCP Cy™5.5 (all BD Biosciences). Cells were distinguished from debris using forward scatter area vs. side scatter area. Doublets were excluded using forward scatter and side scatter (height vs. width). Positive staining thresholds were established using fluorescence minus one (FMO) controls, in which all staining reagents are included except the reagent of interest. For each reagent of interest, the positive gate was set to include 1% of FMO control cells. The cells stained with the reagent of interest that fell within this gate were used to calculate the number of positive cells. Antibody details listed in Table S4.

### Trilineage Differentiation of Parental and Edited WTC Human iPSCs

WTC edited and unedited parental hiPSCs were assessed for pluripotency using STEMdiff™ Trilineage Differentiation Kit (STEMCELL™ Technologies, Inc.) as a means of abbreviated directed differentiation to endoderm (EN), mesoderm (M), and ectoderm (EC), with an undifferentiated (U) control. Cells were cultured according to the STEMdiff™ Trilineage Differentiation Kit protocol in 1:30 diluted Matrigel-coated 24-well plates (Corning). Prior to flow analysis, cells were collected by Accutase detachment, washed in DPBS, and fixed with BD Cytofix™ Fixation Buffer (BD Biosciences) for 30 minutes at 4°C. Cells were washed once with DPBS before pelleting and freezing at -80°C in 50 μL KSR + 10% DMSO. After thawing, cells were washed with 2% BSA in DPBS. EC cells were permeabilized with 0.1% Triton X-100 and 2% BSA in DPBS for 30 minutes at room temperature. EN and M cells were permeabilized with BD Perm/Wash™ buffer (BD Biosciences) for 30 minutes at room temperature. Cells were stained with a lineage-specific antibody (EC: anti-Pax6 AlexaFluor^®^ 647 (BD Biosciences); EN: anti-Sox17 APC (R&D Systems); M: anti-Brachyury APC (R&D Systems) or equal mass of isotype control for 30 minutes at room temperature then washed with their corresponding permeabilization buffer. All cells received a final wash of 2% BSA in DPBS before being resuspended in 2% BSA in DPBS for acquisition. Cells were acquired on a FACSAriaIII Fusion (BD Biosciences) and analyzed using FlowJo V.10.2 (Treestar, Inc.). Samples were gated to exclude debris and cell doublets at previously described. A gate containing 1% of isotype control-positive cells defined the positive staining threshold. Antibody details listed in Table S4.

### In Vitro Directed Differentiation of hiPSCs to Cardiomyocytes

We followed previously reported methods for cardiomyocyte differentiation [60]. Briefly, cells were seeded onto Matrigel-coated 6-well tissue culture plates at a density ranging from 0.75-2×10^6^ cells per well in mTeSR1 supplemented with 1% P/S, 10 μM ROCK inhibitor, and 1 μM CHIR99021 (Cayman Chemical). The following day (designated day 0), directed cardiac differentiation was initiated by treating the cultures with 100 ng/mL ActivinA (R&D Systems) in RPMI media (Invitrogen) containing 1:60 diluted GFR Matrigel (Corning), and insulin-free B27 supplement (Invitrogen). After 17 hours (day 1), cultures were treated with 5 ng/mL BMP4 (R&D systems) in RPMI media containing 1 μM CHIR99021 and insulin-free B27 supplement. At day 3, cultures were treated with 1 μM XAV 939 (Tocris Biosciences) in RPMI media supplemented with insulin-free B27 supplement. On day 5, the media was replaced with RPMI media supplemented with insulin-free B27. From day 7 onwards, media was replaced with RPMI media supplemented with B27 with insulin (Invitrogen).

To measure Troponin T expression, cells were harvested as previously described, fixed with 4% PFA in DPBS for 10 minutes at room temperature, washed twice with 5% FBS in DPBS, and incubated in BD Perm/Wash™ buffer containing anti-cardiac Troponin T AlexaFluor^®^ 647 or equal mass of mIgG1, k AF647 isotype control (all BD Biosciences) for 30 minutes at room temperature (Table S4). After staining, cells were washed with BD Perm/Wash™ buffer, then 5% FBS in DPBS and resuspended in 5% FBS in DPBS with DAPI 2μg/mL). Cells were acquired on a FACSAriaIII Fusion (BD Biosciences) and analyzed using FlowJo software V.10.2. (Treestar, Inc.). Nucleated particles were identified as a sharp, condensed peak on a DAPI histogram and were then gated to exclude doublets as previously described. The cardiac Troponin T (cTnT) positive gate was set to include 1% of cells in the isotype control sample.

### G banding Karyotype Analysis

Karyotype analysis was performed by Diagnostic Cytogenetics Inc. (DCI). A minimum of 20 metaphase cells were analyzed per clone.

### RNA-Seq Analysis

Two clonal populations (one at passage 8 and one at passage 14) were sequenced from the WTC parental line. After dissociation of cell cultures with Accutase, 2-3 x 10^6^ cells were pelleted, washed once with DPBS, re-suspended in 350 μL of Qiagen RLT plus lysis buffer, then flash frozen in liquid nitrogen before storage at -80°C. 101 bp paired end libraries were prepared using an Illumina TruSeq Stranded mRNA Library Prep kit. Libraries were sequenced on an Illumina HiSeq 2500 at a depth of 30 million read pairs (Covance). Adapters were trimmed using Cutadapt [32]. Reads were mapped to human genome build GRCh38 (GCA 000001405.15) and NCBI annotations 107 using STAR aligner. Gene level and isoform level transcript abundances were estimated using Cufflinks.

## Acknowledgements

We thank Michelle Baird, Bruce Conklin, Jacob Corn, Gaudenz Danuser, Daphne Dambournet, Mirek Dundr, Lisa Godsel, Lawrence Goldstein, Kathleen Green, Dirk Hockemeyer, Tony Hyman, Boaz Levi, Miguel Vicente-Manzanares, Wallace Marshall, Tom Misteli, Charles Murry, Lil Pabon, Sean Palecek, Ina Poser, Jennifer Lippincott-Schwartz, William Skarnes, Tim Stearns, Annie Truong, Fyodor Urnov, Xiulan Yang, Jessica Young, and Becky Zaunbrecher for many insightful discussions, advice, and reagents. We also thank Susan Bort, Colette DeLizo, Chris Jeans, and Nadiya Shapovalova for technical support and Thao Do for illustration.

The WTC line that we used to create our gene edited cell lines was provided by the Bruce R. Conklin Laboratory at the Gladstone Institutes and UCSF.

We thank the Allen Institute for Cell Science founder, Paul G. Allen, for his vision, encouragement, and support.

## Supplementary information

### Systematic gene tagging using CRISPR/Cas9 in human stem cells to illuminate cell organization

Brock Roberts^*†^, Amanda Haupt^*^, Andrew Tucker, Tanya Grancharova, Joy Arakaki, Margaret A. Fuqua, Angelique Nelson, Caroline Hookway, Susan A. Ludmann, Irina A. Mueller, Ruian Yang, Alan R. Horwitz, Susanne M. Rafelski, and Ruwanthi N. Gunawardane^†^

Allen Institute for Cell Science, 615 Westlake Ave North, Seattle, WA 98109

^†^Address correspondence to: Brock Roberts, Ruwanthi Gunawardane

^*^These authors contributed equally

Contents:

Supplementary Table S1

Supplementary Table S2

Supplementary Table S3

Supplementary Table S4

Supplementary Figure S1

Supplementary Figure S2

Supplementary Figure S3

Supplementary Figure S4

Supplementary Figure S5

Supplementary Figure S6

Supplementary Figure S7

Supplementary Figure S8

Supplementary Figure S9

Supplementary Figure S10

**Table S1.**
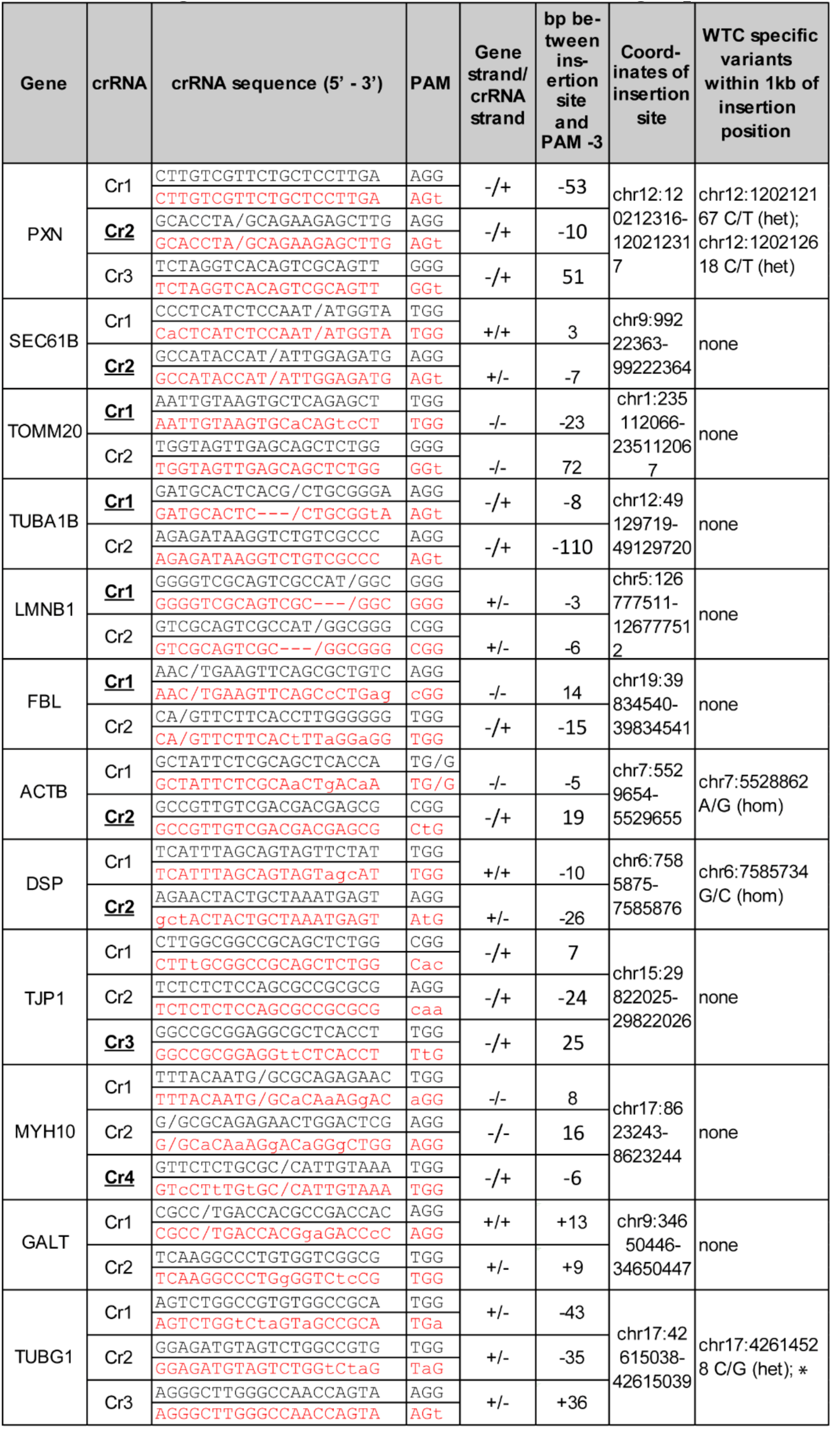
**Design features for crRNAs used in editing experiments shown in Fig. 1D.** Name of the targeted gene, crRNA number, and binding sequence are shown. The crRNA used to create the final clone chosen for expansion and distribution for each gene is bolded and underlined. The non-complementary DNA strand corresponding to the crRNA binding site and PAM in the WTC genome is shown in black. The non-complementary DNA strand corresponding to the crRNA binding site in the donor plasmid and PAM is shown in red. Mutations introduced into the donor plasmid to eliminate Cas9 cleavage are indicated by lower case (point mutations), dashes (deletions), or forward slash (where the tag and linker sequence interrupts the crRNA binding site). The distance between the intended insertion site and the PAM -3 site (where double strand breaks are anticipated) is indicated for each crRNA. Distances are negative when the double strand break is anticipated 5′ of the insertion site and positive when the double strand break is anticipated 3′ of the insertion site relative to gene orientation. Gene orientation and crRNA orientation are defined according to strand in the GRCh38 reference genome. Genomic coordinates are indicated for the site of integration, and for single nucleotide polymorphisms (SNPs) and insertions or deletions (INDELs) specific to the WTC genome within the homology arm region of the plasmid. In cases where the WTC-specific SNP was heterozygous, the reference genome variant was used in the homology arm. Coordinates are from the GRCh38 (GCA 000001405.15) assembly, NCBI annotation 107. *TUBG1 heterozygous SNP was changed to WTC variant in donor plasmid.

**Table S2.** **Clone survival rates after manual isolation.** Number of clones isolated for each enriched population and percentage of clones that survived for initial genetic screening. An asterisk indicates experiments where survival is additionally defined by clones with normal hiPSC morphology. Total percent clones analyzed was calculated for all experiments regardless of whether morphology was a factor considered for survival.

| **Experiment** | **Clones isolated (n)** | **Clones survived (%)** |
| --- | --- | --- |
| SEC61B Cr1 | 53 | 51* |
| SEC61B Cr2 | 65 | 48* |
| FBL Cr1 | 24 | 63* |
| FBL Cr2 | 34 | 15* |
| ACTB Cr1 | 96 | 95 |
| ACTB Cr2 | 96 | 94 |
| TJP1 Cr1 | 45 | 98 |
| TJP1 Cr3 | 48 | 98 |
| LMNB1 Cr1 | 99 | 47* |
| LMNB1 Cr2 | 103 | 45* |
| PXN Cr1 | 24 | 79 |
| PXN Cr2 | 32 | 84 |
| PXN Cr3 | 23 | 96 |
| TOMM20 Cr1 | 96 | 86 |
| DSP Cr1 | 88 | 90 |
| DSP Cr2 | 85 | 100 |
| TUBA1B Cr1 | 96 | 94 |
| TUBA1B Cr2 | 48 | 100 |
| MYH10 Cr1 | 53 | 98 |
| MYH10 Cr2 | 14 | 100 |
| MYH10 Cr4 | 35 | 77 |
| **Total** | **1257** | **79** |

**Table S3.**
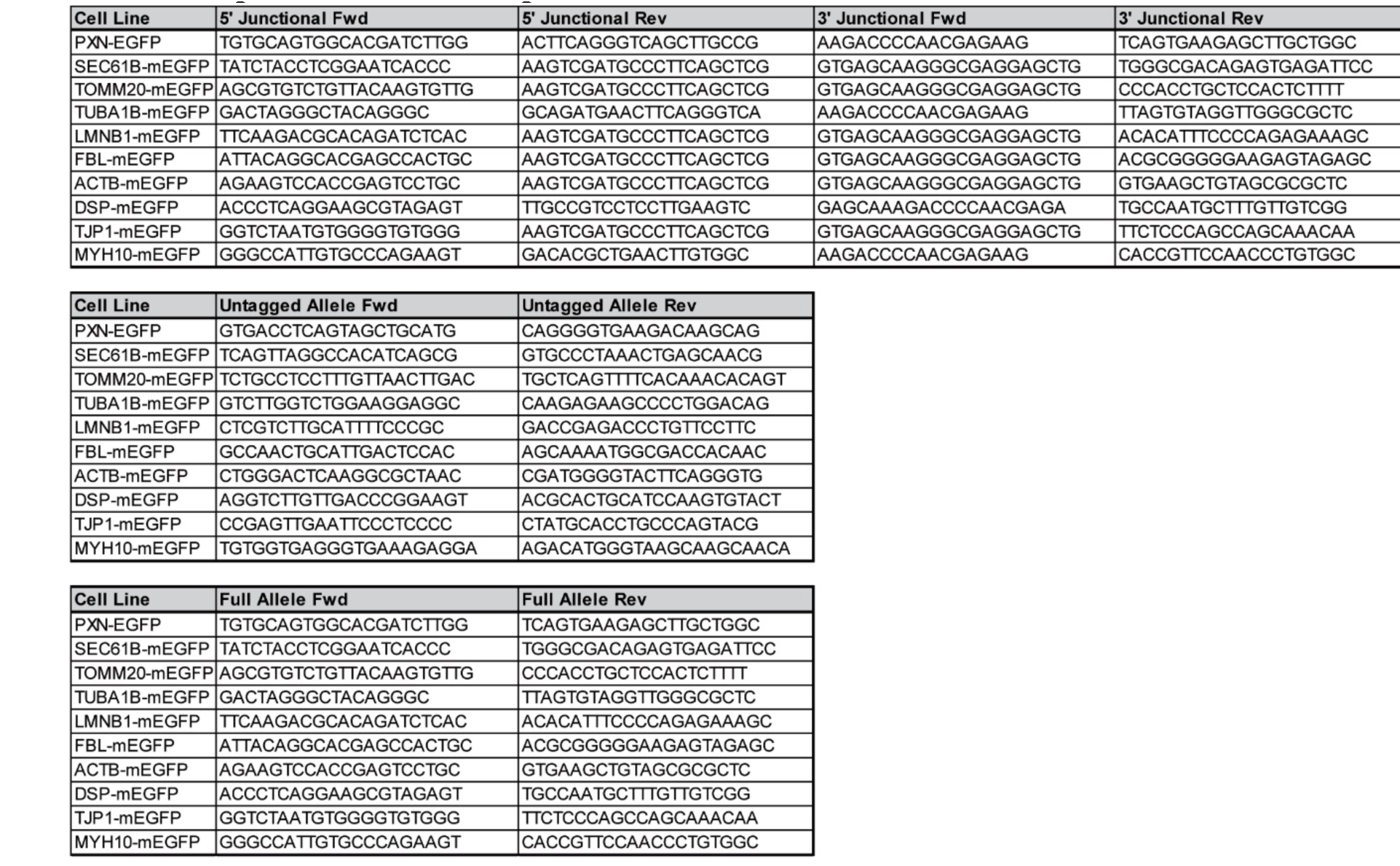
**PCR primers used in experiments.** All primers are listed in 5′ to 3′ orientation.

**Table S4.** **Antibodies used in Western blot, immunofluorescence, and flow cytometry experiments.** Table of antibodies used in Western blots (WB), immunofluorescence (IF), and flow cytometry (Flow) experiments showing dilutions used per application.

| **Primary Antibodies** |  |  |  |  |
| --- | --- | --- | --- | --- |
| **Antibody** | **Type** | **Source** | **Application** | **Secondary Antibody** |
| Alpha tubulin | mouse monoclonal, clone DM1A | ThermoFisher #62204 | WB: 1:10,000<br>IF: 1:250 | goat anti-mouse |
| Beta actin | mouse monoclonal, clone GT5512 | GeneTex #GTX629630 | WB: 1:10,000 | goat anti-mouse |
| Desmoplakin | rabbit polyclonal NW6 | Kathleen Green, Northwestern University | WB: 1:1,000 | goat anti-rabbit |
|  | rabbit polyclonal 1G4 | Kathleen Green, Northwestern University | IF: 1:200 | goat anti-rabbit |
| Fibrillarin | rabbit polyclonal | Abcam #ab5821 | WB: 1:800<br>IF: 1:100 | goat anti-rabbit |
| Lamin B1 (Nuclear lamin B1) | rabbit polyclonal | Abcam #ab16048 | WB: 1:2,000<br>IF: 1:500 | goat anti-rabbit |
| Myosin IIB<br>(Non-muscle myosin heavy chain IIB) | rabbit polyclonal | Cell Signaling Technology #3404 | WB: 1:1,000<br>IF: 1:200 | goat anti-rabbit |
| Paxillin | mouse monoclonal, clone 349 | BD Biosciences #610051 | WB: 1:10,000<br>IF: 1:750 | goat anti-mouse |
| Sec61 beta | rabbit polyclonal | Abcam #ab15576 | WB: 1:10,000<br>IF: 1:250 | goat anti-rabbit |
|  | rabbit polyclonal | Sigma Aldrich #HPA049407 | IF: 1:25 | goat anti-rabbit |
| Tight junction protein ZO1 (ZO1) | rabbit polyclonal | ThermoFisher #617300 | WB: 1:500<br>IF: 1:50 | goat anti-rabbit |
| Tom20 | mouse monoclonal, clone F10 | Santa Cruz Biotechnologies #sc17764 | WB: 1:250<br>IF: 1:100 | goat anti-mouse |
| Beta actin (loading control) | mouse monoclonal, clone BA3R | ThermoFisher #MA515739 | WB: 1:10,000 | goat anti-mouse |
| Alpha actinin (loading control) | mouse monoclonal, clone O.T.02 | ThermoFisher #MA191860 | WB: 1:2,000 | goat anti-mouse |
| GFP | mouse monoclonal mix, clones 7.1 and 13.1 | Sigma Aldrich #11814460001 | WB: 1:250 | goat anti-mouse |
| **Directly Conjugated Primary Antibodies** |  |  |  |  |
| **Antibody** | **Type** | **Source** | **Application** |  |
| TRA-1-60, Brilliant Violet™ 510 | mouse monoclonal, clone TRA-1-60 | BD Biosciences #563188 | Flow: 1:40 |  |
| SSEA-3, Alexa Fluor® 647 | rat monoclonal, clone MC-631 | BD Biosciences #561145 | Flow: 1:160 |  |
| SSEA-1, Brilliant Violet™ 421 | mouse monoclonal, clone MC480 | BD Biosciences #562705 | Flow: 1:2,500 |  |
| Nanog, Alexa Fluor® 647 | mouse monoclonal, clone N31-355 | BD Biosciences #561300 | Flow: 1:67 |  |
| Oct3/4, Brilliant Violet™ 510 | mouse monoclonal, clone 40/Oct-3 | BD Biosciences #563524 | Flow: 1:67 |  |
| Sox2, V450 | mouse monoclonal, clone O30-678 | BD Biosciences #561610 | Flow: 1:80 |  |
| TRA-1-60, PerCP-Cy™5.5 | mouse monoclonal, clone TRA-1-60 | BD Biosciences #561573 | Flow: 1:25 |  |
| SSEA-3, PE | rat monoclonal, clone MC-631 | BD Biosciences #560237 | Flow: 1:10 |  |
| SSEA-1, Brilliant Violet™ 650 | mouse monoclonal, clone H198 | BD Biosciences #564232 | Flow: 1:10 |  |
| Oct3/4, PerCP-Cy™5.5 | mouse monoclonal, clone 40/Oct-3 | BD Biosciences #560794 | Flow: 1:5 |  |
| Sox2, Alexa Fluor® 647 | mouse monoclonal, clone 245610 | BD Biosciences #560294 | Flow: 1:25 |  |
| Nanog, PE | mouse monoclonal, clone N31-355 | BD Biosciences #560483 | Flow: 1:2.5 |  |
| Cardiac Troponin T, Alexa Fluor® 647 | mouse monoclonal, clone 13-11 | BD Biosciences #565744 | Flow: 1:200<br>IF: 1:200 |  |
| Mouse IgG1 k Isotype Control, Alexa Fluor® 647 | mouse monoclonal, clone MOPC-21 | BD Biosciences #565571 | Flow: 1:200 |  |
| Brachyury, APC | goat polyclonal | R&D Systems #IC2085A | Flow: 1:40 |  |
| Goat IgG Isotype Control, APC | goat polyclonal | R&D Systems #IC108A | Flow: 1:40 |  |
| Sox17, APC | goat polyclonal | R&D Systems #IC1924A | Flow: 1:80 |  |
| Goat IgG Isotype Control, APC | goat polyclonal | R&D Systems #IC108A | Flow: 1:80 |  |
| Pax6, Alexa Fluor® 647 | mouse monoclonal, clone O18-1330 | BD Biosciences #562249 | Flow: 1:20 |  |
| Mouse IgG2a k Isotype Control, Alexa Fluor® 647 | mouse monoclonal, clone MOPC-173 | BD Biosciences #558053 | Flow: 1:10 |  |
| **Secondary Antibodies** |  |  |  |  |
| **Antibody** | **Type** | **Source** | **Application** |  |
| goat anti-mouse IgG (H+L),<br>Alexa Fluor® 647 conjugate | goat polyclonal | ThermoFisher #A21236 | WB: 1:10,000<br>IF: 1:500 |  |
| goat anti-rabbit IgG (H+L),<br>Alexa Fluor® 647 conjugate | goat polyclonal | ThermoFisher #A21245 | WB: 1:10,000<br>IF: 1:500 |  |

**Figure S1.**
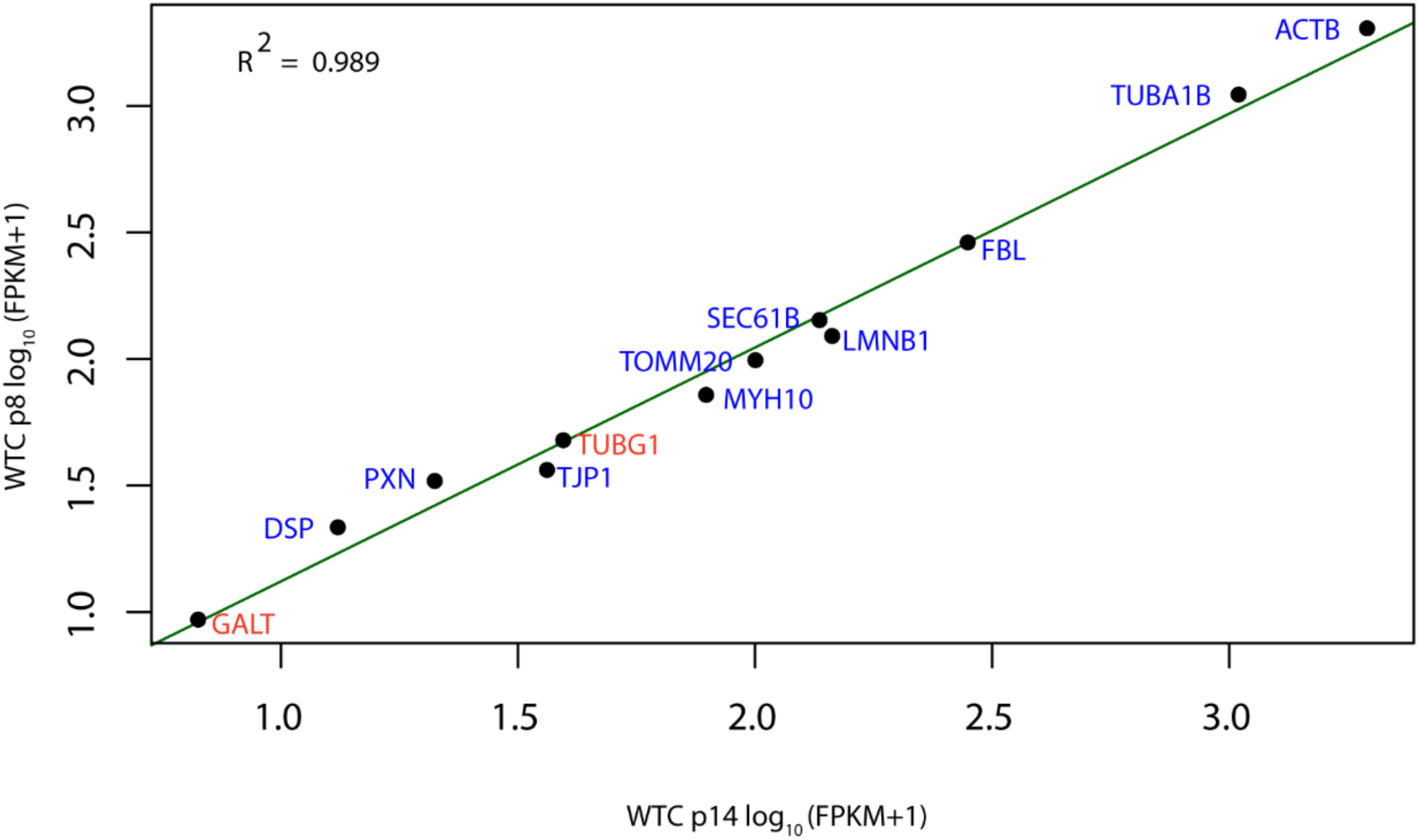
Expression levels of the 12 genes attempted for genome editing in the WTC parental cell line. Transcript abundance for each gene was estimated from RNA-Seq data. Samples were derived from the WTC parental line after 8 passages (p8) in culture and 14 passages (p14) in culture, as indicated. Transcript abundances were calculated in units of fragments per kilobase of transcript per million fragments mapped (FPKM). Log_10_ (FPKM+1) transcript abundances from parental WTC p8 and p14 samples were plotted against each other and were highly correlated (R^2^= 0.989). The two genes (TUBG1 and GALT) that were not successfully edited (Fig. S3) are highlighted in red.

**Figure S2.**
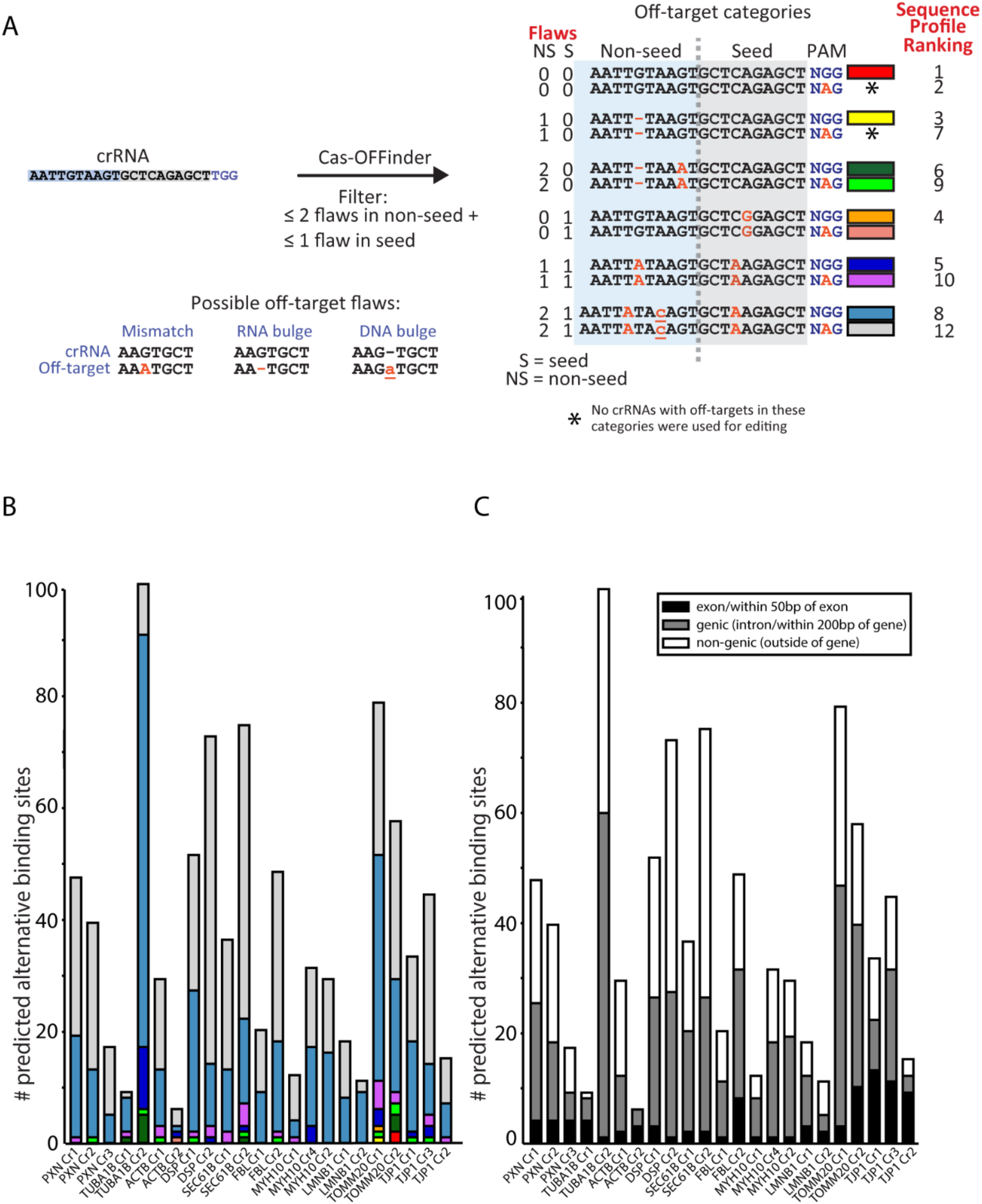
Predicted genome wide CRISPR/Cas9 alternative binding sites, categorized according to sequence profile and location with respect to genes. (**A**) Predicted alternative CRISPR/Cas9 binding sites are categorized for each crRNA used. Each predicted off-target sequence was categorized according to its sequence profile (the number of mismatches and RNA or DNA bulges it contains relative to the crRNA used in the experiment) and position relative to the PAM. Cas-OFFinder was used to identify all alternative sites genome-wide with ≤2 mismatches/bulges in the non-seed and/or ≤1 mismatch/bulge in the seed region, with an NGG or NAG PAM. As indicated, the seed and non-seed region of a crRNA binding sequence was defined with respect to its proximity to the PAM sequence. Overlapping Cas-OFFinder results with the same double strand break site were collapsed into one category using sequence profile ranking. (**B**) Predicted off-target sequence breakdown based on sequence profile (colors refer to categories defined in (A). (**C**) All predicted off-target sites were additionally categorized according to their location with respect to annotated genes. Genomic location was defined as follows; <u>exon</u>: inside exon or within 50 bp of exon; <u>genic</u>: in intron (but >50 bp from an exon) or within 200 bp of an annotated gene; <u>non-genic</u>: >200 bp from an annotated gene. See methods for more details on off-target categorization.

**Figure S3.**
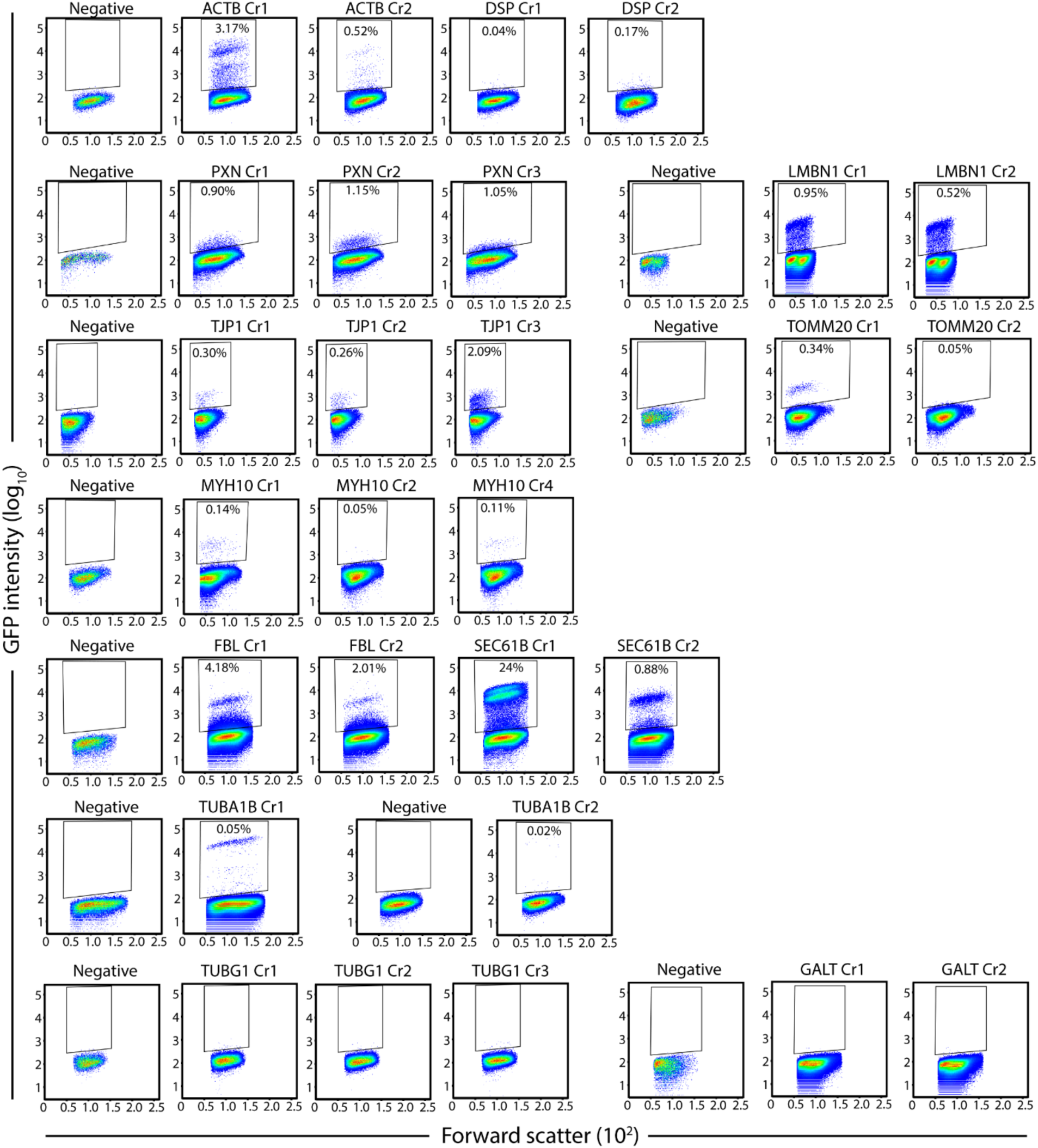
Flow cytometry plots of edited cells post-transfection. Data from all experiments are shown, as in Fig. 1C, along with the untransfected negative control accompanying each experiment. Gates (boxes) indicate the population of putatively edited cells and values reflect the percentage of edited cells within the total population, except for unedited cells (negative) and GALT and TUBG1 experiments where enrichment was not possible (bottom panel). Forward scatter is shown on the x-axis. Data are grouped and displayed with the negative control performed in that experiment.

**Figure S4.**
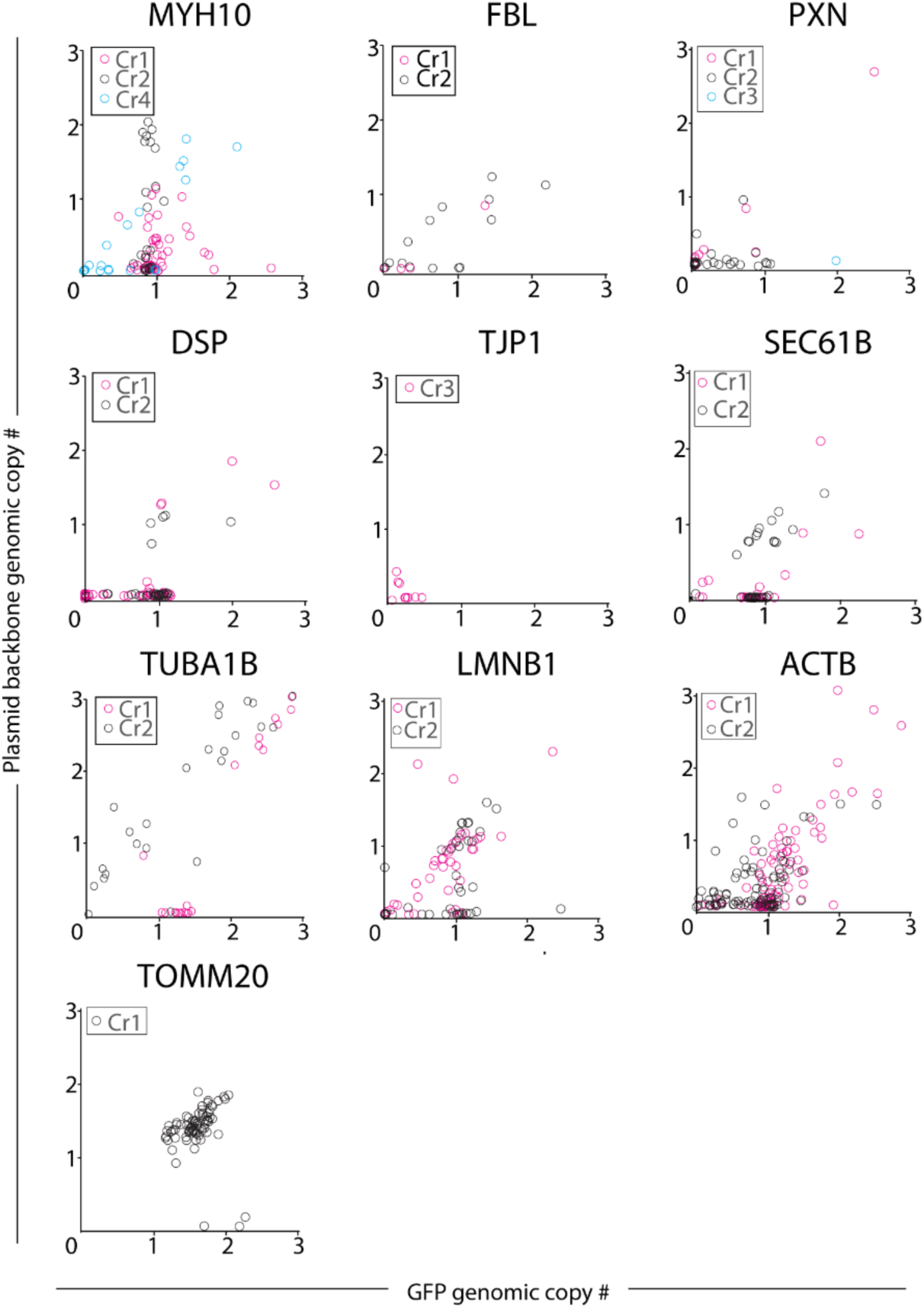
ddPCR screening data. ddPCR screening data for all experiments (Fig. 2A, step 1). Each data point represents one clone. Clones with GFP genomic copy number of ∼1 to ∼2 and plasmid backbone genomic copy number <0.2 were typically considered for further analysis. TJP1 clones consistently produced GFP copy number values <1 despite validation by junctional PCR, imaging and Western blot as putative mono-allelic clones. This result is unresolved and under investigation.

**Figure S5.**
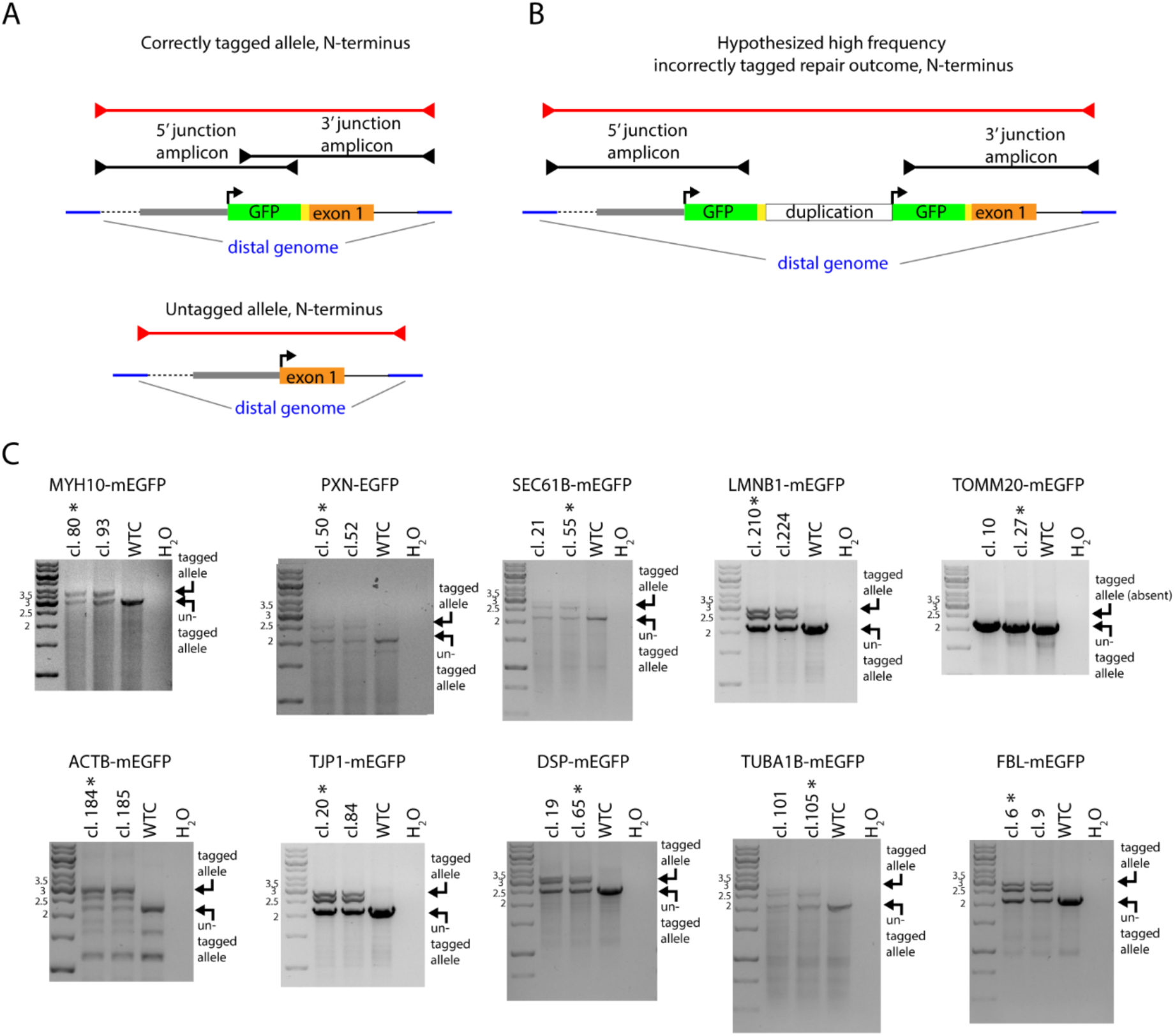
Amplification of complete junctional (non-tiled) PCR products to demonstrate presence of the allele anticipated from tiled junctional PCR product data. (A) Junctional PCR primers complementary to sequences flanking the homology arms in the distal genome (also used in tiled junctional PCR assays, shown in black), were used together to co-amplify tagged and untagged alleles (red). N-terminal tag shown as an example. (B) This assay served to rule out anticipated DNA repair outcomes where tiled junctional PCR data leads to a misleading result because the GFP tag sequence has been duplicated during HDR, as indicated by the schematic. An N-terminal tag duplication is shown as an example. (C) Molecular weight markers are as indicated (kb). Two final clones (indicated by “cl. #”) are represented for each experiment. Asterisk indicates the final clone chosen for distribution and imaging. A band intermediate in size between the anticipated tagged and untagged allele products is consistently observed, which we hypothesized corresponds to a heteroduplex of the tagged and untagged allele products.

**Figure S6.**
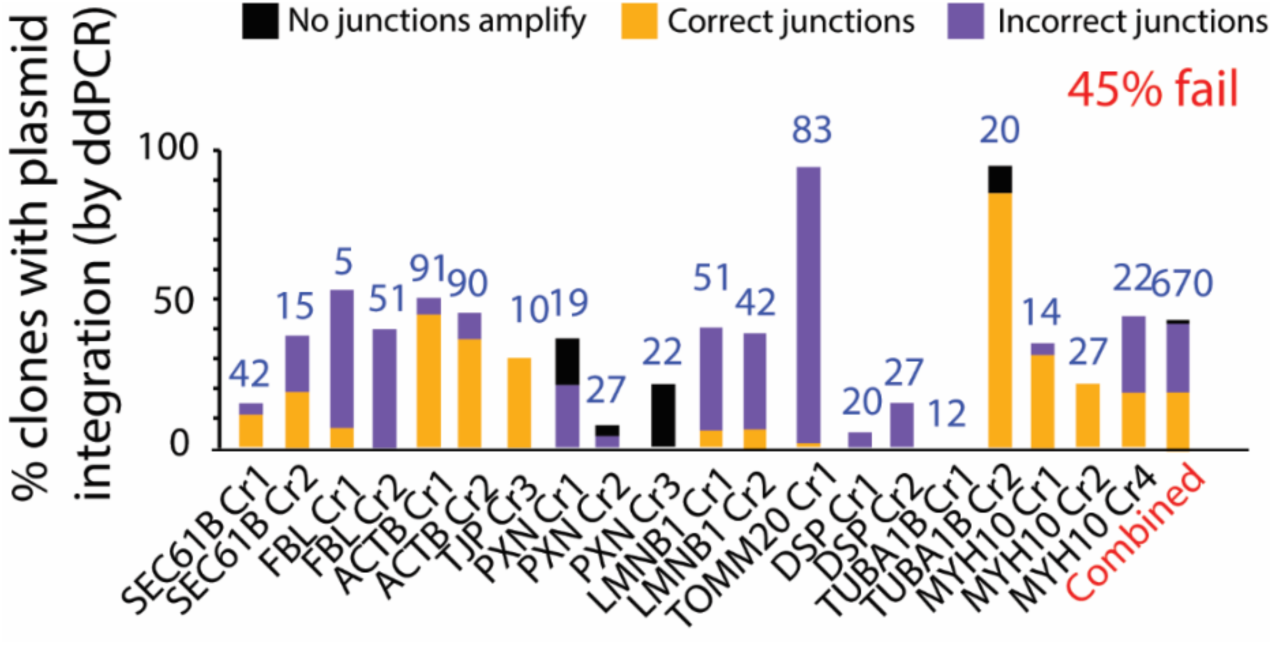
Junctional PCR analysis of clones rejected by ddPCR due to integrated plasmid. The percentage of clones in each experiment with KAN/AMP copy number ≥0.2 is displayed on the y-axis. Stacked bars represent the 3 subcategories of clones observed within clones rejected according to this criterion. In clones where at least one incorrect junction is amplified (purple), the aberrant junction product is larger than expected. These cases are interpreted as possible evidence of plasmid backbone integration at the targeted locus. Clones in which no junctions are amplified (black) are interpreted to contain random integration of the donor plasmid, because the absence of the junction product indicates that the plasmid has no physical association with the target locus. These cases may alternatively represent high levels of transfected plasmid persisting in cells. Clones in which both junctions are correct (yellow) are interpreted to contain duplications of the GFP tag sequence at the targeted locus. As shown in Fig. 2, amplification of the correct junctional product occurs in 90% of clones confirmed by ddPCR to lack plasmid integration while also containing the GFP tag. By contrast, the majority of clones rejected by ddPCR (45% of total) due to suspected integrated donor plasmid yield incorrect junctional products. We hypothesize that integration of the donor plasmid tends to cause incorrect junctional products and that plasmid incorporation at the site of intended editing is a common byproduct of HDR. We note that plasmid incorporation at the site of intended editing may result in duplications of the tag sequence and thus junction amplicons with the anticipated size may be frequently observed in clones where the plasmid backbone incorporates at the target locus (Fig. S5).

**Figure S7.**
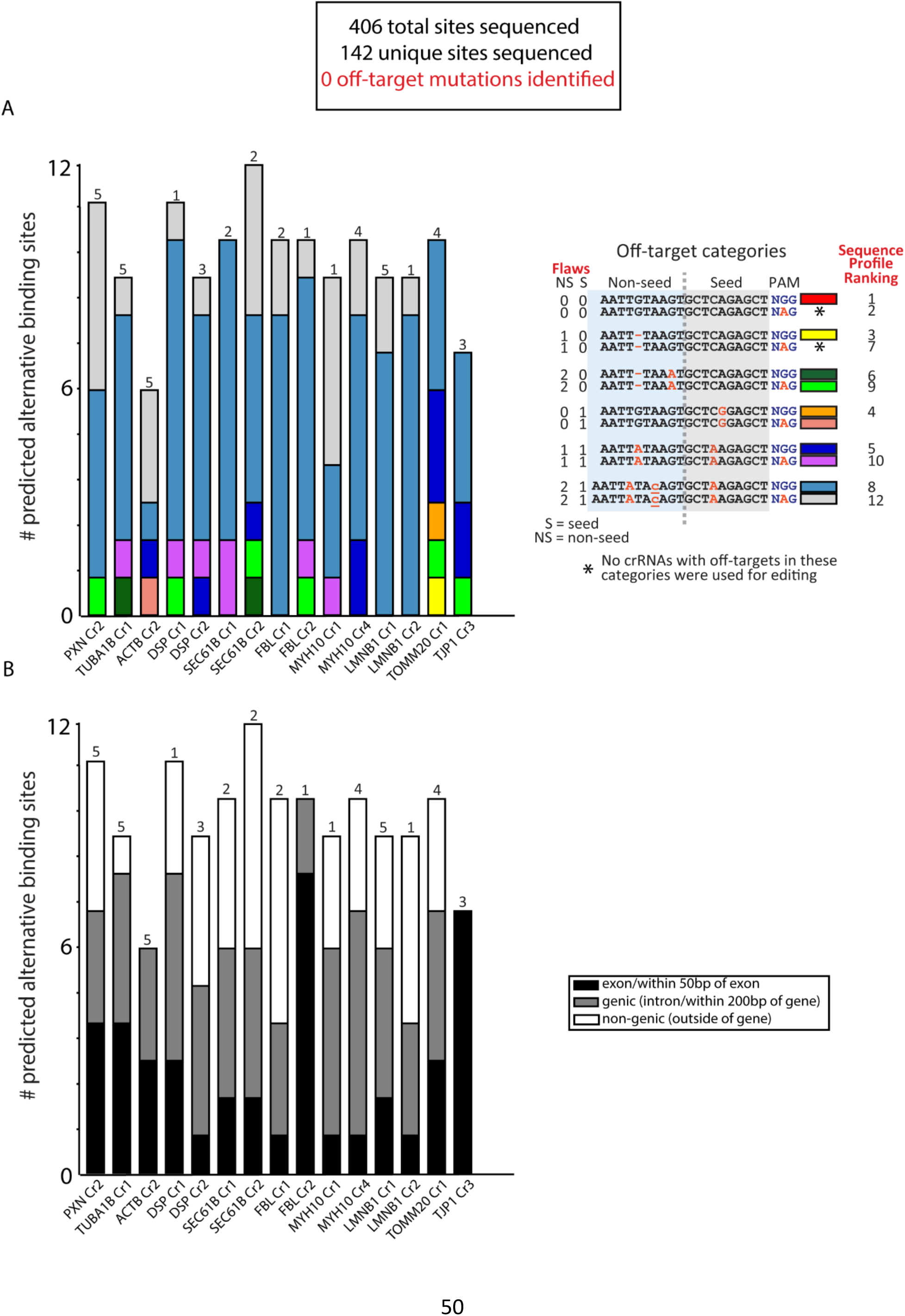
Sequenced CRISPR/Cas9 alternative binding sites, categorized according to sequence profile and location with respect to genes. A subset of CRISPR/Cas9 alternative binding sites identified by Cas-OFFinder (Fig. S2) were selected for sequencing. Numbers above bars represent the number of clones sequenced for each experiment. All 406 sequenced sites were found to be wild type. (**A**) Breakdown of sequenced off-target sites by sequence profile. Overlapping Cas-OFFinder results with the same double strand break site were collapsed into one category using sequence profile ranking. (**B**) Breakdown of sequenced off-target sites by genomic location with respect to annotated genes. Genomic location was defined as follows; <u>exon</u>: inside exon or ≤50 bp of exon; <u>genic</u>: in intron (but >50 bp from an exon) or ≤ 200 bp of an annotated gene; <u>non-genic</u>: >200 bp from an annotated gene. See methods for more details on off-target categorization.

**Figure S8.**
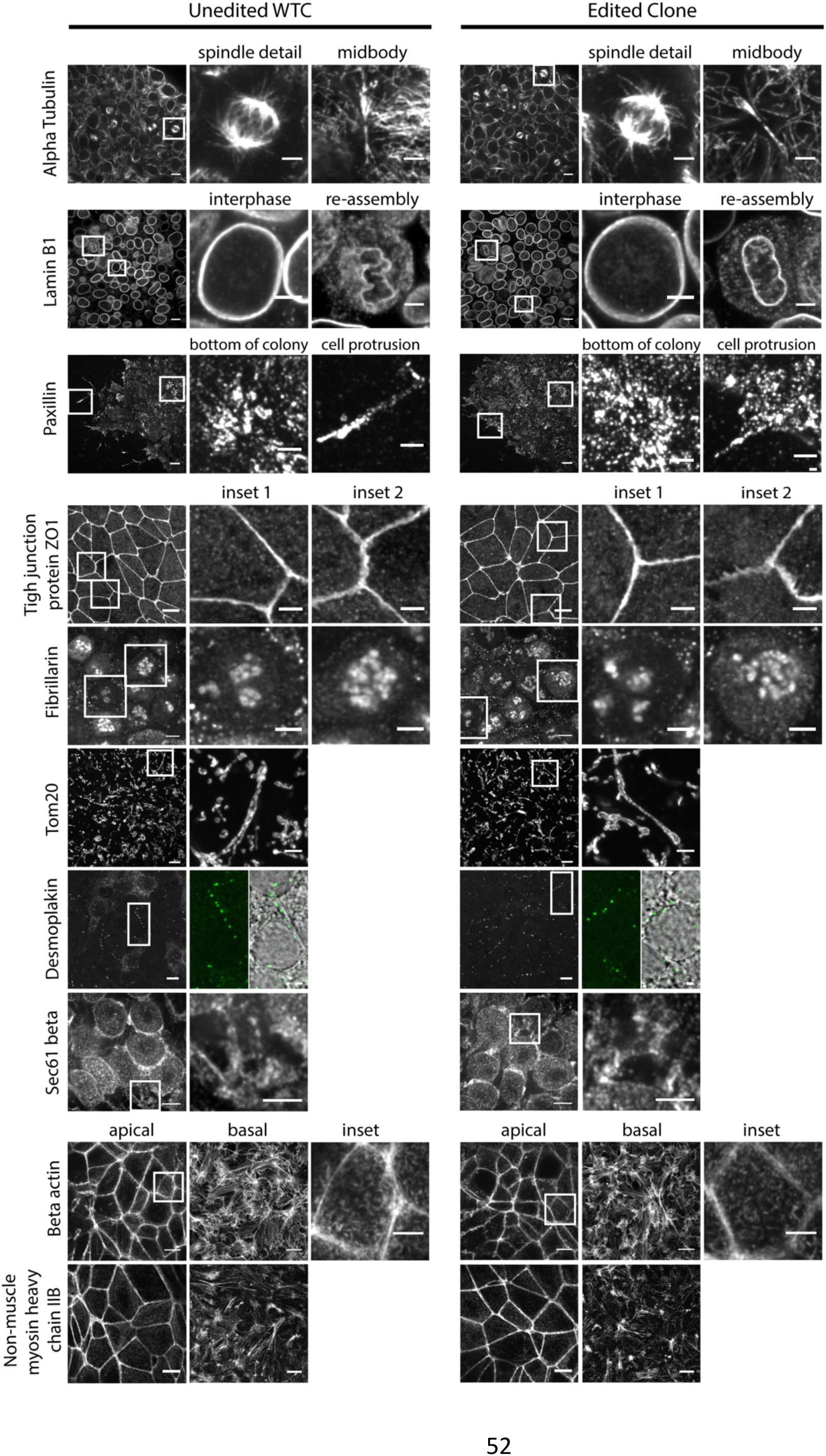
Comparison of unedited versus edited cells by immunofluorescence. Labeled structures in unedited WTC parental cells and edited cell lines are compared. Whole field of views (FOVs) shown on the left, with insets highlighted with white boxes. **Alpha tubulin** panel: anti-alpha tubulin antibody staining FOV with insets highlighting a spindle and a midbody (midbody inset was obtained from an apical image slice, not shown in the FOV). Images represent single z-section slices. FOV scale bar is 10 μm, insets are 3 μm. **Lamin B1** panel: anti-lamin B1 antibody staining FOV with insets illustrating interphase and nuclear envelope reassembly. Images represent maximum intensity projections of 3 apical z-sections. FOV scale bar is 10 μm, insets are 3 μm. **Paxillin** panel: anti-paxillin antibody staining FOV with insets highlighting the basal cell surface and cell protrusions in detail. Images represent maximum intensity projections of 3 basal z-sections. FOV scale bar is 10 μm, insets are 3 μm. **Tight junction protein ZO1** panel: anti-ZO1 antibody staining FOV with two insets. Images represent maximum intensity projections of 10 apical z-sections. FOV scale bar is 10 μm, insets are 3 μm. **Fibrillarin** panel: anti-fibrillarin antibody staining FOV with two insets illustrating variation in nucleolar staining. Images represent a single apical z-section. FOV scale bar is 5 μm, insets are 3 μm. **Tom20** panel: anti-Tom20 antibody staining FOV with one inset highlighting a single mitochondrial tubule. Images represent maximum intensity projections of 4 basal z-sections. FOV scale bar is 10 μm, inset is 3 μm. **Desmoplakin** panel: anti-desmoplakin staining FOV with one inset showing the GFP channel and transmitted light image overlay to show desmoplakin puncta localization at the cell-cell boundaries. Images represent maximum intensity projections of z-sections spanning the entire colony and single z plane for the transmitted light image. FOV scale bar is 10 μm, inset is 1 μm. Sec61 beta panel: anti-Sec61 beta antibody staining FOV with one inset. Images represent maximum intensity projections of 3 z-sections near the middle of the cell colony. FOV scale bar is 8 μm, inset is 4 μm. **Beta actin** panel: Phalloidin-Rhodamine staining showing apical and basal FOVs, and an apical region inset. Images represent maximum intensity projections of either apical or basal z-sections. Apical and basal image scale bars are 10 μm, inset is 4 μm. **Non-muscle myosin heavy chain IIB** panel: anti-myosin IIB antibody staining FOV showing apical and basal regions. Images represent maximum intensity projections of 4 apical or basal z-sections of the cell colony. Scale bars are 10 μm. All images acquired on a spinning disk confocal microscope except panels shown for desmoplakin, which was acquired on a laser scanning confocal microscope. Antibody and method details are available in Table S4.http://www.allencell.org/

**Figure S9.**
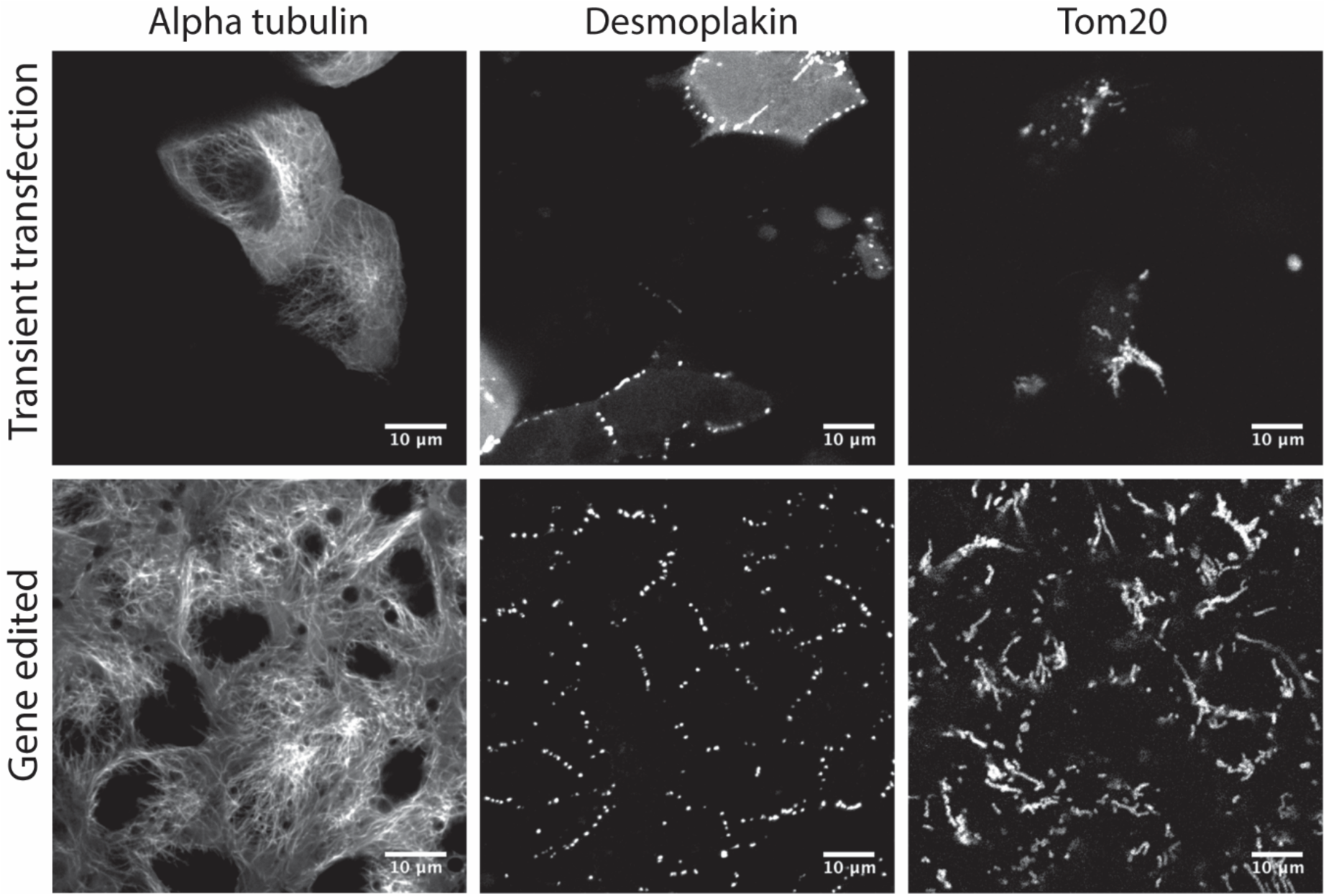
Live cell imaging comparison of transiently transfected cells and genome edited cells. Top panels depict transiently transfected WTC cells and lower panels depict gene edited clonal lines. **Left**: WTC transfected with mEGFP-tagged alpha tubulin construct compared to the TUBA1B-mEGFP edited cell line. Images are a single apical frame. **Middle**: WTC transfected with EGFP-tagged desmoplakin construct compared to the DSP-mEGFP edited cell line. Images are maximum intensity projections of apical 4 z-frames. **Right**: WTC transfected with mCherry-tagged Tom20 construct compared to the TOMM20-mEGFP edited cell line. Images are single basal frames of the cell. All imaging was performed in 3D on live cells using laser-scanning confocal microscope. Movie versions of these z-stacks can be found at http://www.allencell.org/.

**Figure S10.**
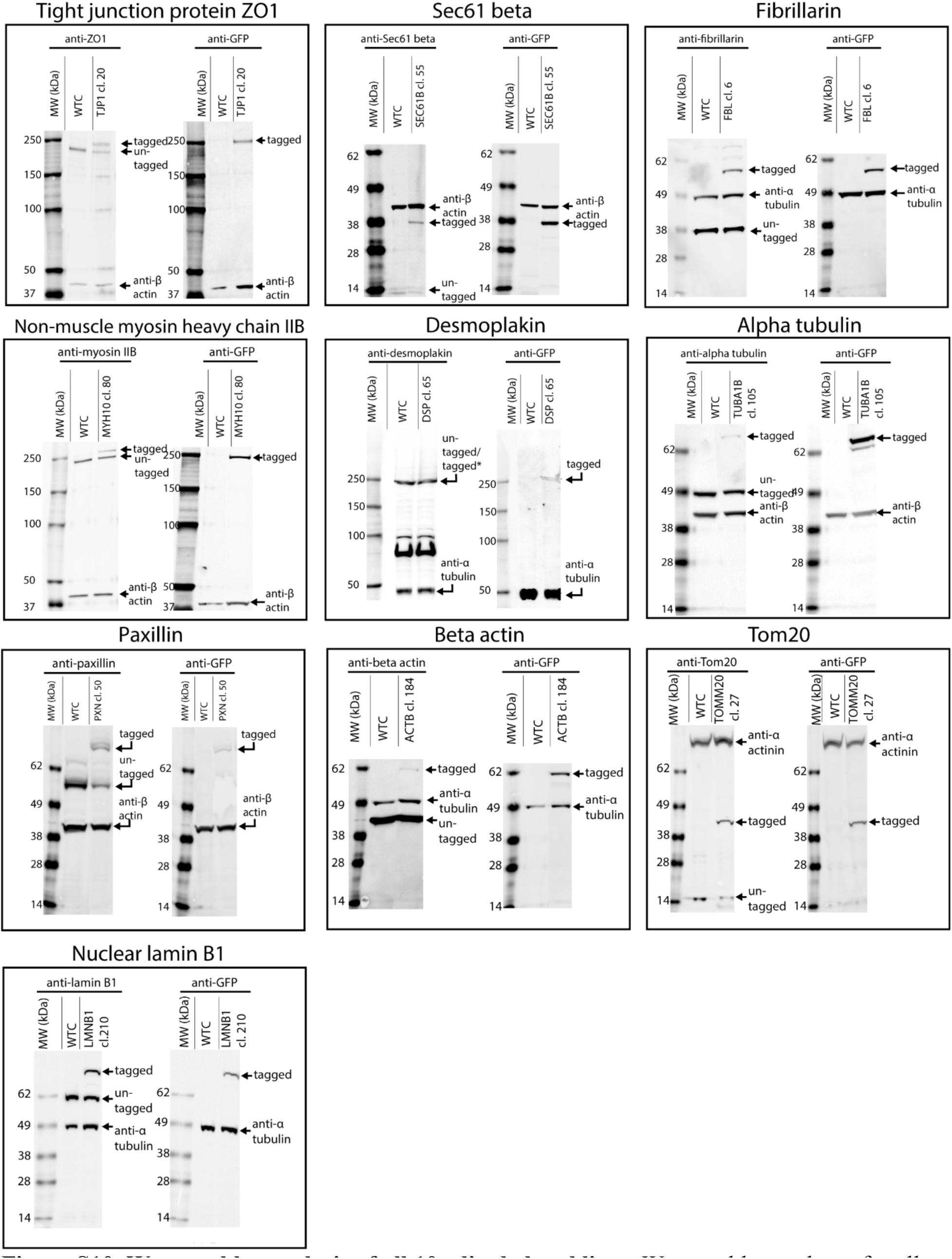
Western blot analysis of all 10 edited clonal lines. Western blot analyses for all experiments are presented as in Fig. 4. Proteins and antibodies used on different blots are as indicated. In all cases, blots with antibodies against the respective target proteins are shown in the left blot and show the tagged and untagged protein products. Separation of untagged and tagged protein versions from the mEGFP-tagged desmoplakin clone was not possible due to the large size of the target protein (asterisk). Blots with anti-GFP antibodies showing only the tagged protein are shown in the right blot, as indicated. Alpha actinin, beta actin, and alpha tubulin were used as loading controls, as indicated. Lysates from unedited cells and the edited clonal lines are as indicated, as are bands corresponding to the labeled, predicted proteins. Antibody information is available in Table S4.

